# Disruption of prepulse inhibition is associated with severity of compulsive behavior and nucleus accumbens dopamine receptor changes in Sapap3 knockout mice

**DOI:** 10.1101/694935

**Authors:** Elizabeth E Manning, Abigail Y Wang, Linda M Saikali, Anna S Winner, Susanne E Ahmari

## Abstract

Obsessive compulsive disorder (OCD) is associated with disruption of sensorimotor gating, which may contribute to difficulties inhibiting intrusive thoughts and compulsive rituals. Neural mechanisms underlying these disturbances are unclear; however, striatal dopamine is implicated in regulation of sensorimotor gating and OCD pathophysiology. The goal of this study was to examine the relationships between sensorimotor gating, compulsive behavior, and striatal dopamine receptor levels in Sapap3 knockout mice (KOs), a widely used preclinical model system for OCD research. We found a trend for disruption of sensorimotor gating in Sapap3-KOs using the translational measure prepulse inhibition (PPI); however, there was significant heterogeneity in both PPI and compulsive grooming in KOs. Disruption of PPI was significantly correlated with a more severe compulsive phenotype. In addition, PPI disruption and compulsive grooming severity were associated with reduced dopamine D1 and D2/3 receptor density in the nucleus accumbens core (NAcC). Compulsive grooming progressively worsened in Sapap3-KOs tested longitudinally, but PPI disruption was first detected in high-grooming KOs at 7 months of age. Through detailed characterization of individual differences in OCD-relevant behavioral and neurochemical measures, our findings suggest that NAcC dopamine receptor changes may be involved in disruption of sensorimotor gating and compulsive behavior relevant to OCD.

## Introduction

Obsessive compulsive disorder (OCD) is a severe neuropsychiatric disorder affecting 1-3% of the population^1-3^. Intrusive thoughts (obsessions) and uncontrollable actions (compulsions) are the core clinical features of OCD. Subjects with OCD also show disturbances in sensorimotor gating^4-8^, a filtering process that impacts the content and organization of what is consciously perceived by an individual. It has been proposed that sensorimotor gating deficits in OCD may facilitate intrusive thoughts and impair inhibitory control over compulsive actions, although these relationships have not been clearly demonstrated in patients.

Underlying mechanisms responsible for disruption of sensorimotor gating in OCD are unclear; however, disturbances in striatal dopamine have been implicated in both the regulation of sensorimotor gating and manifestation of compulsive behavior. In rodent models sensorimotor gating is disrupted by increased dopamine neurotransmission following systemic administration of dopamine agonists/releasers^9,10^ or site specific manipulations in dorsal or ventral striatum^11-13^. Dopamine system disturbances have also been described in the striatum in patients with OCD^14-18^ using *in vivo* molecular imaging with single photon emission computerized tomography (SPECT) and positron emission tomography (PET). Binding of striatal dopamine D1 and D2/3 receptors is reduced in OCD patients^14-16,19^, and dopamine transporter binding is increased^17,20^. It has been proposed that these alterations may represent compensatory changes in response to elevated dopamine tone, consistent with evidence that transdiagnostic compulsive behaviors are associated with elevated striatal dopamine. Specifically, increased cue-evoked striatal dopamine release has been observed via PET imaging in cocaine users, subjects with binge eating disorder, and patients with Parkinson’s disease who develop compulsive behavior following dopamine agonist treatment^21-24^. There is also evidence that drugs that increase dopamine signalling can exacerbate compulsive behavior in OCD patients^25^. Further supporting the role of elevated dopamine in the manifestation of OCD symptoms, dopamine antagonists can be useful as an adjunct therapy in OCD patients that don’t show clinical improvement following treatment with first-line selective serotonin reuptake inhibitors (SSRIs)^26^. Together, these results suggest a potential relationship between alterations in striatal dopamine signalling and disruption of both sensorimotor gating and compulsive behavior in OCD.

Preclinical model systems can be used to gain mechanistic insight into the relationship between dopamine signalling, impaired sensorimotor gating, and other OCD-relevant behaviors using a straightforward translational paradigm: prepulse inhibition (PPI) of the acoustic startle response^27^. For example, studies in mouse models suggest that elevated striatal dopamine and disruption of dorsal striatum function may have relevance to disruption of PPI in Tourette Syndrome^28,29^. Although prior studies have demonstrated PPI deficits in pharmacologic models of relevance to OCD^30,31^, sensorimotor gating deficits and their relationship to compulsive behaviors have not been characterized in transgenic mouse models. Here, we examined PPI in Sapap3 knockout mice (KOs), a widely used model system for OCD research^32-34^ that displays compulsive grooming, anxiety^32^, and cognitive changes in tasks examining reversal learning^35,36^ and habit learning^37,38^. Sapap3-KOs also show hyperactivity in the dorsal and central striatum (CS)^32-34,39^ that would be predicted to influence PPI; in contrast, the ventral striatum has not been thoroughly characterized in this model system^40^. The goal of this study was therefore to probe the relationship between PPI, striatal dopamine receptor levels, and compulsive behavior in Sapap3-KOs using behavioral testing and *ex vivo* autoradiography.

## Results

### Disruption of PPI is associated with compulsive grooming phenotype in Sapap3-KOs

PPI was examined in male Sapap3-KO and wild-type (WT) control mice at ∼8 months of age, when approximately half of KOs (10/18) showed lesions resulting from compulsive grooming. Analysis of average %PPI revealed a trend for differences between genotypes (Figure 1A; unpaired Welch’s T-test: p=0.10, t_27.28_=1.70), with both groups showing similar increased PPI with higher prepulse intensities (Figure S1A; main effect prepulse, F_(2,56)_= 87.4, p<0.0001). However, visual inspection of the data and an F test to compare variance between the groups demonstrated more variability in KOs compared to WTs (F_(17,11)_=3.3, p=0.0499), prompting us to determine whether this heterogeneity within KOs mapped onto other behavioral measures. As expected, Sapap3-KOs groomed significantly more than WTs (Figure 1B; Mann-Whitney test: WT: M=38.5, KO: M=628, p<0.0001). In addition, we determined that a subgroup of Sapap3-KOs with skin lesions (KO-L, n=10, unfilled symbols Figure 1B) showed elevated grooming relative to KOs without lesions (KO-NL, n=8, filled symbols), supporting the idea that lesions result from elevated grooming. However, KO-NL also groomed significantly more than WTs (Bonferroni post-hoc test, p=0.012; also see Figure S2F-H for data from an independent cohort).

**Figure 1:**
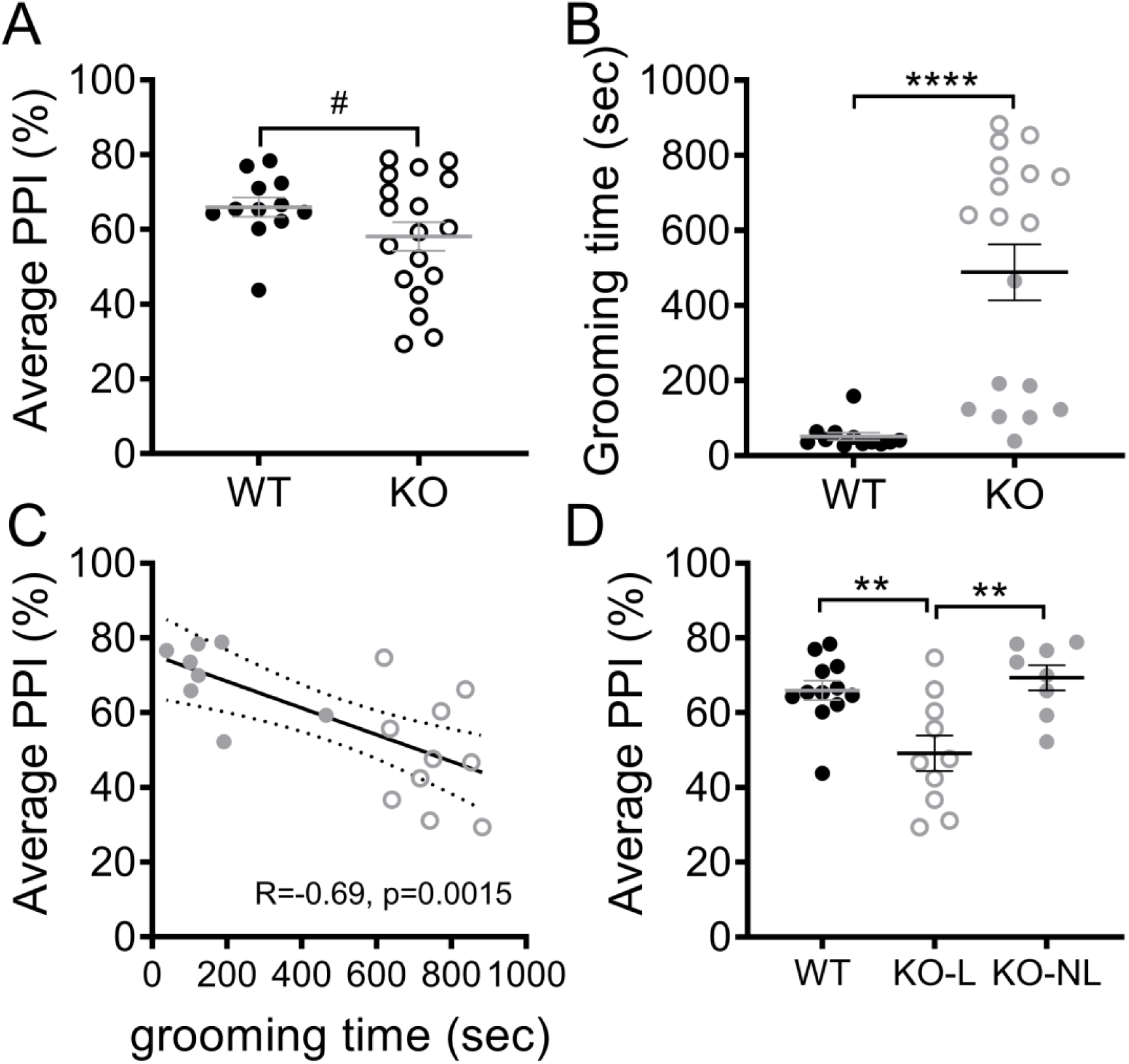
Disruption of PPI is associated with compulsive grooming phenotype in Sapap3-KOs. **A)** PPI (average % across 3 prepulse intensities tested) did not differ between Sapap3-KO and WT mice. **B)** Time spent grooming was significantly elevated in Sapap3-KO relative to WT, as previously described. Unfilled grey symbols denote KO-L, filled grey symbols denote KO-NL. **C)** Average PPI was significantly negatively correlated with time spent grooming, suggesting disruption of PPI in Sapap3-KOs with the most severe compulsive grooming phenotype. **D)** When Sapap3-KOs were separated based on the presence (KO-L) or absence (KO-NL) of lesions resulting from compulsive grooming, significant disruption of PPI in KO-L was revealed relative to WT and KO-NL (Bonferroni post-hoc tests: WT vs KO-L, p=0.006; WT vs KO-NL, p>0.99, KO-L vs KO-NL p=0.003). In panel A, # indicates results of unpaired Welch’s T test, in panel B * indicates results of Mann-Whitney test, in panel D * indicate results of Bonferroni post-hoc test. **** p<0.0001; ** p<0.01; # p=0.10. n= 12 WT, 18 KO (10 KO-L, 8 KO-NL). PPI = prepulse inhibition, KO= knockout, KO-L= KOs with lesions, KO-NL= KOs without lesions, WT= wild-type.

Given the significant heterogeneity in compulsive grooming phenotype in Sapap3-KOs, the relationship between PPI and grooming levels was assessed to determine whether PPI deficits were observed in mice with more severe compulsive grooming. In Sapap3-KOs, average %PPI was significantly negatively correlated with time spent grooming (Figure 1C; KO: R=-0.69, p=0.0015; WT: R=-0.10, p=0.75), and was selectively impaired in KO-L relative to WTs and KO-NL (Figure 1D; main effect of group, F_(2,27)_=8.5, p=0.0013). Importantly, startle amplitude during blocks 2 and 3, which is used to calculate %PPI, did not differ between groups (main effect of group, p=0.43; Figure S1B).

### Individual differences in compulsive grooming and disruption of PPI in Sapap3-KOs are associated with changes in striatal dopamine receptor binding density

To determine whether heterogeneous disruption of PPI and grooming severity in Sapap3-KOs were associated with changes in striatal dopamine receptor density, *ex vivo* autoradiography was used to measure binding of D1 and D2/3 in CS and ventral striatum (nucleus accumbens core; NAcC) of animals described in Figure 1 (see Figure 2A for regions of interest). Receptor densities were then correlated with behavioral measures in Sapap3-KOs. D2/3 receptor density was significantly different between the genotypes (Figure 2B; main effect genotype, F_(1,25)_=4.6, p=0.042), and there was a trend for interaction with striatal region of interest suggesting that this effect may be more prominent in NAcC (genotype x region interaction, p=0.097). Reduced D2/3 density in KOs was also associated with individual differences in compulsive grooming and disruption of PPI. Again this effect was selective to NAcC, with reduction of D2/3 binding associated with both disruption of PPI (Figure 2C, R=0.59, p=0.017) and increased grooming (Figure 2D, R=-0.68, p=0.004). In contrast, D2/3 density in CS showed no association with these OCD-relevant behaviors (Figure 2E-F; PPI: p=0.46, R=0.20; grooming: p=0.72, R=-0.01).

**Figure 2.**
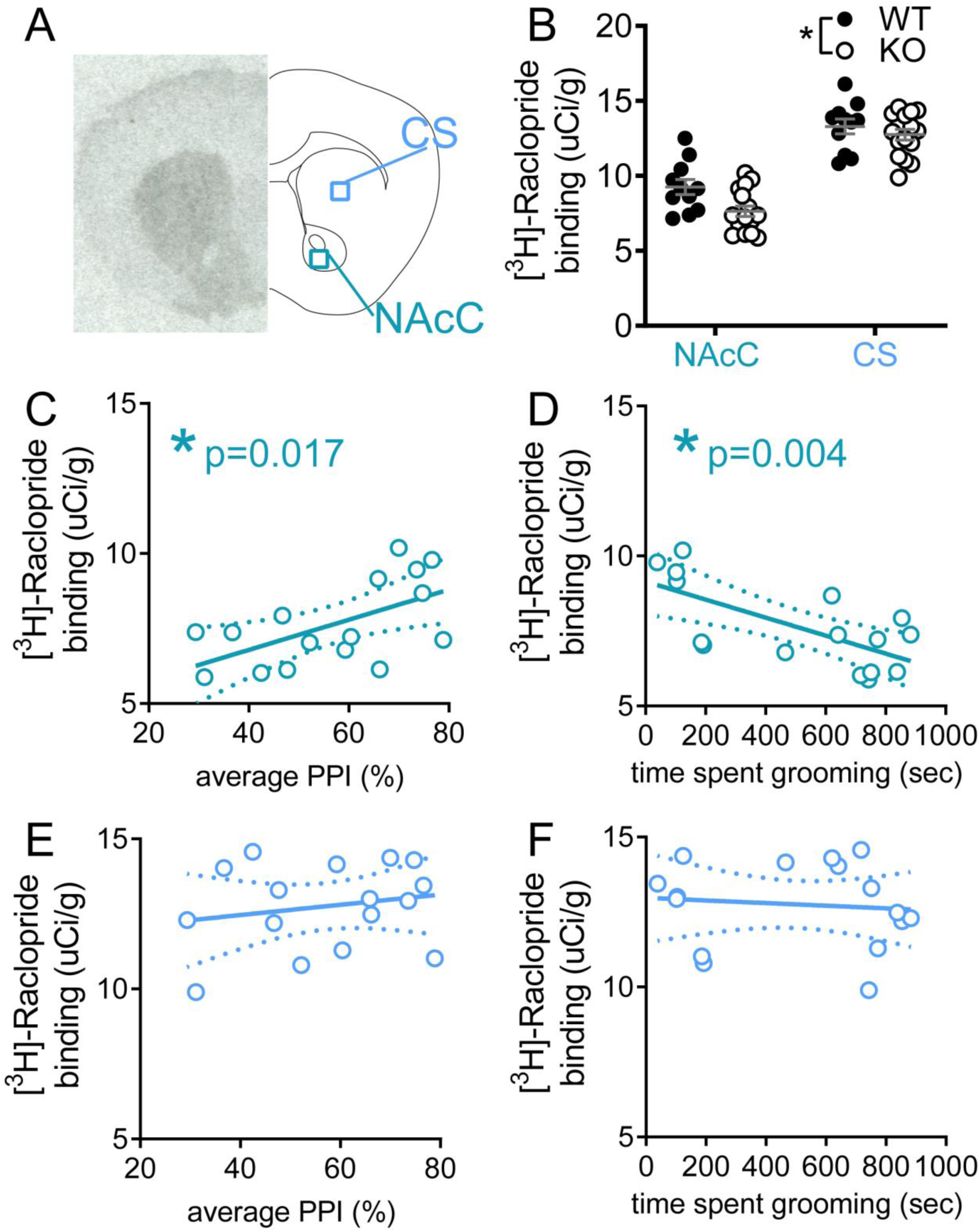
Reduced dopamine D2/3 receptor density in NAcC is associated with severity of OCD-relevant phenotypes in Sapap3-KO mice. **A)** Autoradiography was performed in striatal tissue to quantify density of D2/3 receptors in NAcC and CS using [^3^H]-Raclopride. (Left) Representative autoradiograph for total binding. (Right) Schematic of coronal brain section. **B)** D2/3 was significantly reduced in Sapap3-KO. Reduced D2/3 binding in NAcC was associated with lower %PPI **(C)** and increased severity of compulsive grooming **(D)** in Sapap3-KOs. D2/3 density in CS showed no association with %PPI **(E)** or grooming **(F)** in Sapap3-KOs. In panel B, * indicates significant difference between WT and KOs (p<0.05), in panels C-D * indicates significant correlation (p<0.05). n= 11 WT, 16 KO. PPI = prepulse inhibition, CS= central striatum, NAcC= Nucleus accumbens core, KO= Sapap3 knockout, WT= wild-type.

D1 receptor density was not significantly different between genotypes (Figure 3B, p=0.74); however, reduced D1 receptor density was associated with increased severity of OCD-relevant behaviors in Sapap3-KO mice. Specifically, in the NAcC, disruption of PPI was significantly correlated with reduced D1 density (Figure 3C, NAcC: R=0.54, p=0.021), and there was a trend for correlation between increased grooming and reduced D1 density (Figure 3D, NAcC: R=-0.43, p=0.074). In CS, there was also a trend for correlation between disruption of PPI and reduced D1 density (Figure 3E, CS: R=0.40, p=0.096), but D1 density was not associated with individual differences in grooming severity (Figure 3F, R=-0.35, p=0.16).

**Figure 3.**
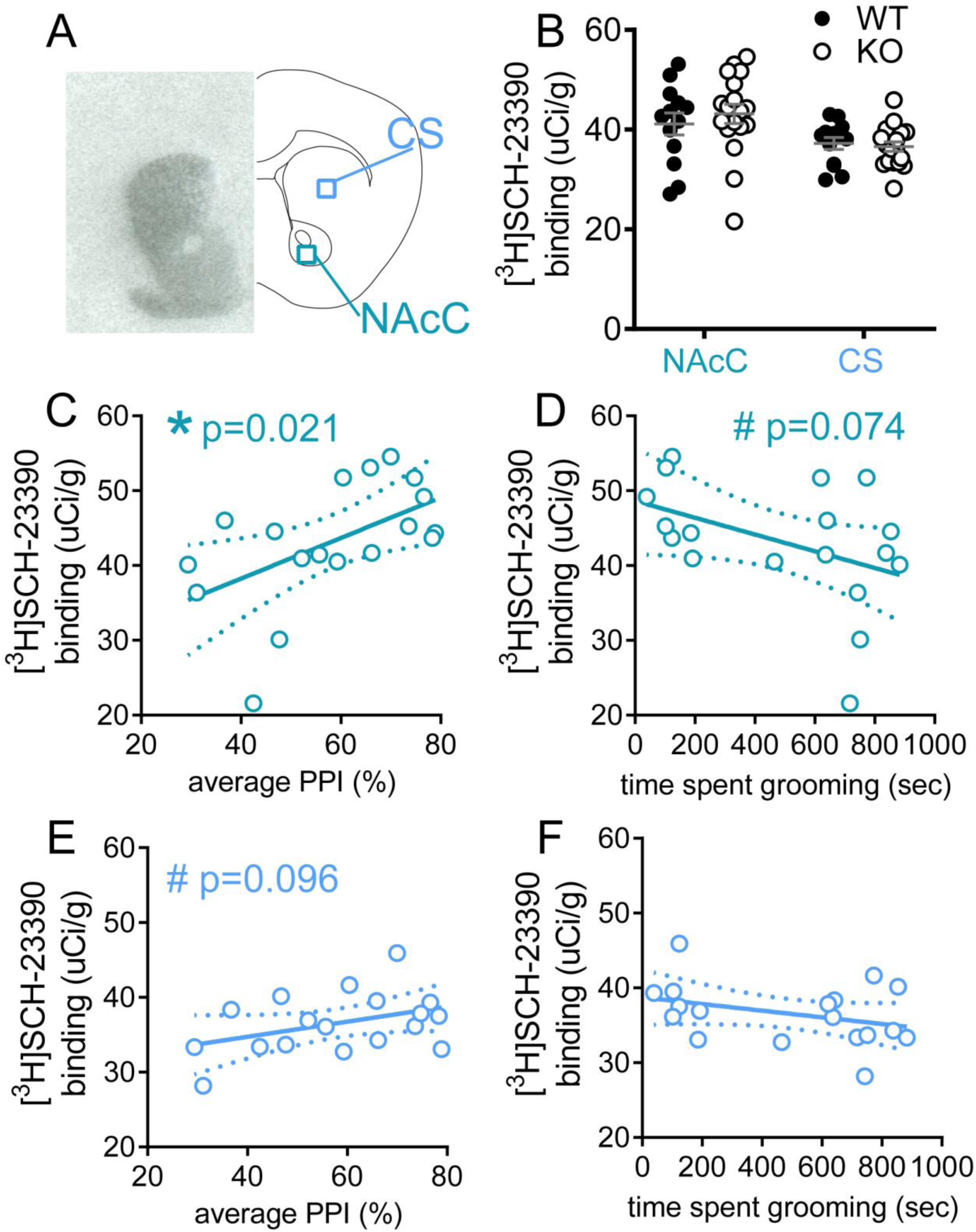
Reduced dopamine D1 receptor density in NAcC is associated with severity of disruption of PPI in Sapap3-KO mice. **A)** Autoradiography was performed in striatal tissue to quantify density of D1 receptors in CS and NAcC using [^3^H]-SCH-23390. Representative autoradiograph for total binding is shown (left) and schematic of coronal brain section (right). **B)** D1 density was unchanged in Sapap3-KOs. In Sapap3-KOs, reduced D1 in NAcC was significantly correlated with reduced PPI **(C)** and showed a trend association with severity of compulsive grooming **(D)**. D1 density in CS showed a trend correlation with PPI **(E)** and no association with compulsive grooming in Sapap3-KOs **(F)**. * indicates significant correlation, # indicates trend correlation (0.05<p<0.10). n= 13 WT, 18 KO. PPI = prepulse inhibition, CS= central striatum, NAcC= Nucleus accumbens core, KO= Sapap3 knockout, WT= wild-type.

### Dopamine transporter binding density is increased in striatum of Sapap3-KO mice

Striatal dopamine system changes were also examined by measuring binding density for the dopamine transporter (DAT) using ex vivo autoradiography. DAT density was increased in Sapap3-KOs compared to WT littermates (Figure 4B, main effect of genotype, F_(1,29)_=4.8; p=0.037). There was a trend for genotype x striatal subregion interaction (p=0.094), suggesting that this effect may be more prominent in CS. In contrast to striatal dopamine receptor densities, DAT binding density in NAcC and CS was not correlated with changes in PPI or grooming in Sapap3-KOs (NAcC: Figure 4C-D, PPI p=0.36, grooming p=0.52; CS: Figure 4E-F; PPI p=0.71, grooming p=0.62).

**Figure 4.**
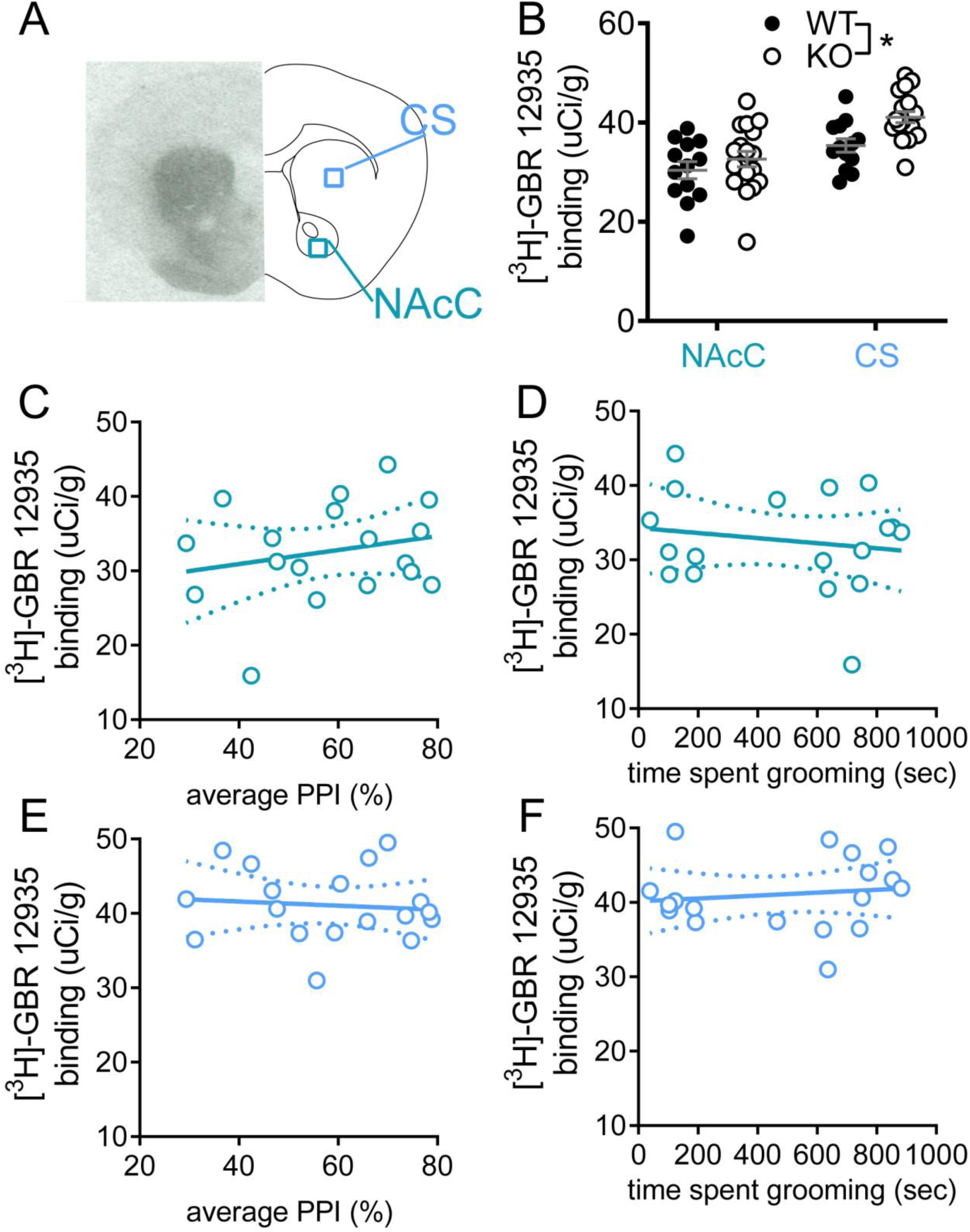
DAT binding in striatum is increased in Sapap3-KOs. **A)** Autoradiography was performed in striatal tissue to quantify density of DAT in CS and NAcC using [^3^H]-GBR-12935. Representative autoradiograph for total binding is shown in the left panel, with schematic of coronal brain section on the right. **B)** DAT density was elevated in Sapap3-KOs. * indicates significant difference between WT and KO (p<0.05). DAT binding in NACc was not correlated with disruption of PPI **(C)** or grooming severity **(D)** in Sapap3-KOs. DAT binding in CS showed no association with PPI **(E)** or grooming **(F)** in Sapap3-KOs. n= 13 WT, 18 KO. PPI = prepulse inhibition, CS= central striatum, NAcC= Nucleus accumbens core, DAT= dopamine transporter, KO= Sapap3 knockout, WT= wild-type.

### Comparison of the developmental progression of compulsive grooming, lesions, and PPI impairment

It is hypothesized that impaired sensorimotor gating in OCD may contribute to disruption of inhibitory control over compulsive actions. To test whether disruption of PPI in Sapap3-KOs might precede development of the compulsive grooming phenotype, and thereby contribute to the onset of compulsive behavior, PPI and grooming were both assessed monthly from 2 months (before the development of lesions) until 8 months of age (when approximately half of KOs have lesions) in male and female Sapap3-KOs and WT littermates. Time spent grooming progressively increased with age (Figure 5A, genotype x age interaction: F_(6,270)_=15.8, p<0.0001), with significantly elevated grooming detected at 5 months in Sapap3-KOs (Bonferroni post-hoc tests). The number of Sapap3-KOs with lesions resulting from severe compulsive grooming increased with age (Figure 2SF-H; 6 months: n=5; 7 months: n=9, 8 months; n=12). Note, in other cohorts tested weekly in our laboratory at an earlier age, a modest elevation in grooming was detected relative to WT littermates as early as 7 weeks of age (Figure S3 and ^39^); however, this small effect-size difference was not detectable during the less frequent (monthly) testing performed in the current study.

**Figure 5.**
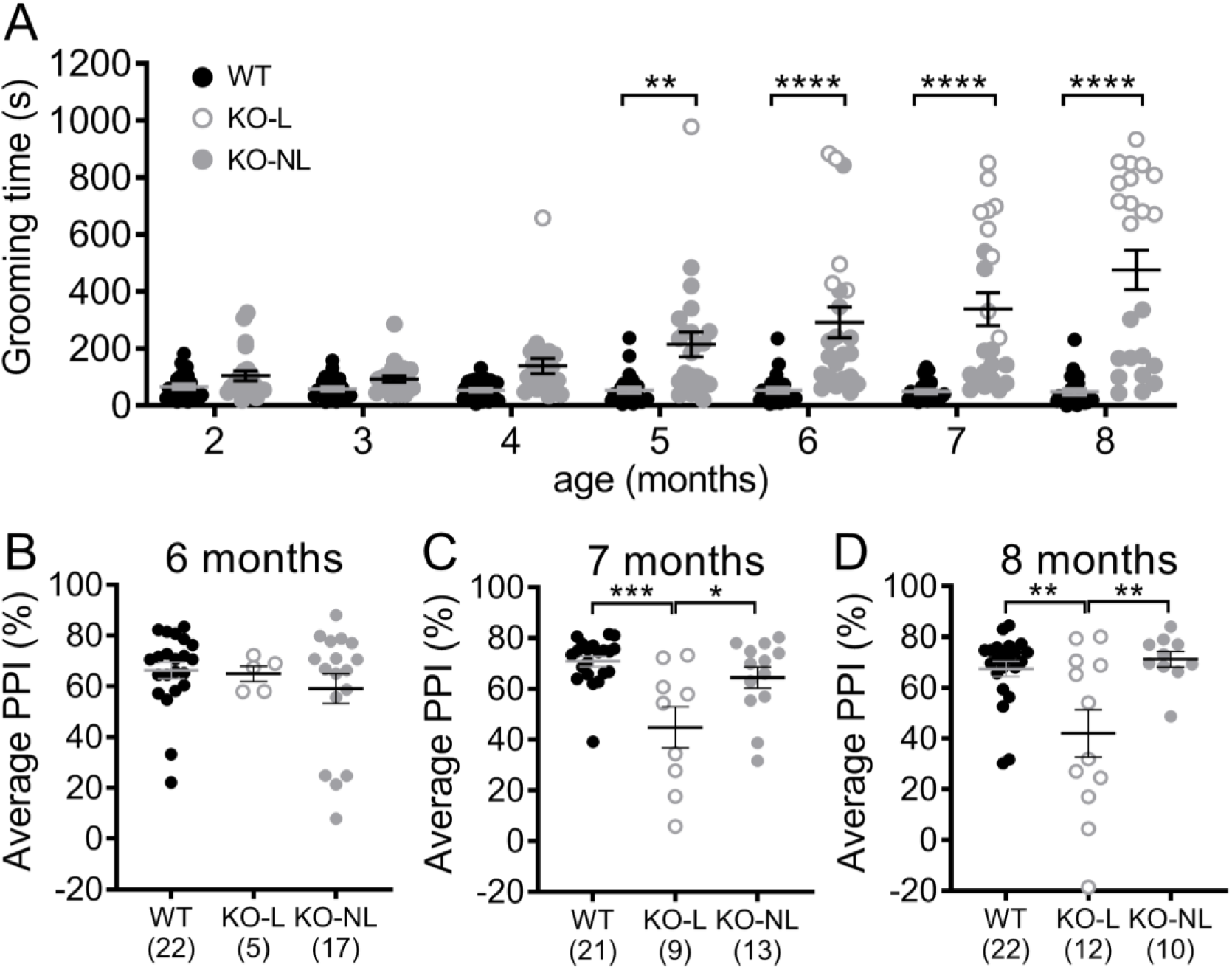
Comparison of the developmental progression of excessive grooming, lesions, and PPI impairment. A) Longitudinal progression of compulsive grooming phenotype in Sapap3-KOs from 2-8 months of age. Unfilled grey symbols are KO-L; grey filled symbols are KO-NL. Grooming was significantly increased in Sapap3-KOs relative to WT from 5 months of age. B-D). PPI was compared between groups at 6-8 months of age when startle did not significantly differ between genotypes (see Figure S2B). Although there are no group differences at 6 months of age, PPI disruption emerges in KO-L at 7 months of age. * indicates significant Bonferroni post-hoc test comparing KO to WT (panel A) or KO-L to comparison group (panel B-D). **** p<0.0001, ** p<0.01, * p<0.05. n= 24 WT, 23 KO (panel A). KO= Sapap3 knockout, KO-L= KOs with lesions, KO-NL= KOs without lesions, WT= wild-type, PPI= prepulse inhibition.

When comparing PPI between WTs and KOs across development (not taking grooming severity into account), there was a trend towards reduced PPI in KOs (Figure S2A, p=0.095), but no interaction with age (p=0.93) (note: PPI increased with age consistent with previous reports^41,42^; main effect of age: F_(6,270)_=29.2, p<0.0001). However, analysis of startle amplitude revealed age-dependent genotype differences (Figure S2B, genotype x age interaction F_(6,270)_=3.8, p=0.001). Whereas startle amplitude decreased in WT mice as they aged (consistent with other studies in mice^41,42^), KOs had relatively stable startle amplitude over time. Since group differences in startle amplitude confound interpretation of PPI, we therefore focused analysis on 6-8 months of age when startle amplitude did not differ between the groups (Figure S2C-E). PPI was compared between KO-L, KO-NL and WT to account for effects of compulsive grooming phenotype. This demonstrated that PPI was not significantly different between groups at 6 months (Figure 5B, F_(2,41)_=0.7, p=0.49); however, KO-L showed reduced PPI relative to the other two groups at 7 and 8 months (Figure 5C-D, 7 months: F_(2,40)_=9.2, p=0.0005; 8 months: F_(2,41)_=7.8, p=0.0013).

## Discussion

Here, we demonstrate that Sapap3-KOs show impaired sensorimotor gating as measured by disruption of PPI, but found that these deficits only manifested when mice progressed to a severe compulsive grooming phenotype. Autoradiography revealed that disruption of PPI was associated with reduced D1 and D2/3 receptor binding in the NAcC. In addition, severity of compulsive grooming was associated with reduced D2/3 binding in NAcC. Finally, DAT binding was elevated in KOs independent of the striatal region examined. Our findings are consistent with PET/SPECT imaging in patients with OCD, and may reflect compensatory responses to striatal hyperdopaminergic tone that are more prominent in the NAcC. Together, these observations suggest that alterations in dopamine signalling within the NAcC may contribute to OCD-relevant behavioral disturbances in Sapap3-KOs, and that longer exposure to altered dopamine signalling may be necessary to impact PPI.

The observed association between severity of compulsive grooming and disruption of PPI in the Sapap3-KO model could reflect that one of these behaviors plays a causal role in the presentation of the other, or that a common neural mechanism(s) contributes to the development or manifestation of both behaviors. A clear relationship between PPI and other OCD symptoms has not been established in previous clinical studies. Early studies in medicated patients demonstrated that severity of PPI impairment was significantly associated with severity of OCD symptoms as measured by the Yale-Brown Obsessive Compulsive Scale (Y-BOCS)^7,8^. Although these associations have not been replicated in more recent studies examining unmedicated OCD patients^4,6^, reduced PPI in patients with tics was noted in one of these samples, supporting the hypothesis that disruption of PPI may be associated with lack of inhibitory control over repetitive actions^4,5^. Future studies seeking to examine the relationship between OCD symptoms and other behavioral measures will likely benefit from a large sample that reflects the broad range of patient severity present across the diagnosis of OCD, similar to our current examination of PPI in Sapap3-KOs displaying a range of severity of compulsive behavior.

Interestingly, we recently demonstrated that disruption of cognitive flexibility in a reversal learning task is not associated with severity of compulsive grooming in Sapap3-KOs^36^, and these findings have been replicated by other laboratories^35,43^. Others have also recently demonstrated aberrant habit learning in Sapap3-KOs, and have begun to examine potential relationships between compulsive grooming and flexible cognition in this context^37,38^. In the first study, increased habitual behavior in Sapap3-KOs (insensitivity to outcome devaluation) was not directly compared to compulsive grooming^38^. However, mice were 7-12 weeks old at the start of habit training, suggesting that a severe compulsive grooming phenotype was not present based on the results of our study. In the second study, which used different testing conditions that specifically promoted habitual behavior in WT mice, Sapap3-KOs showed more goal-directed behavior (sensitive to outcome devaluation). Although analysis of the relationship between habitual behavior and compulsive grooming was underpowered, there was a trend for a bias towards habitual behavior in low grooming Sapap3-KOs and flexible goal-directed behavior in high grooming Sapap3-KOs^37^. These studies of cognitive flexibility largely support the idea that OCD-relevant cognitive changes and compulsive grooming are distinct behavioral domains in Sapap3-KOs, whereas our current findings suggest that compulsive grooming and disruption of PPI in Sapap3-KOs may be related. The Sapap3-KO model therefore serves as a useful system for further dissection of the contributions of overlapping versus distinct circuits associated with different OCD-relevant behavioral domains. This is an important question given that distinct patterns of activity in striatum and prefrontal cortex are associated with symptoms vs cognitive deficits^44^. Better understanding of these behavior-specific neural changes may thus be valuable for guiding the development of more effective treatments.

Striatal dopamine signalling is implicated in both the pathophysiology of OCD and regulation of PPI. Our autoradiography results are consistent with PET/SPECT imaging studies demonstrating reduced D2/3 and D1 binding in striatum of OCD patients^15,16,19^. Compulsive behavior and disruption of PPI are both typically associated with a hyperdopaminergic state in striatum^13,21-25,45^. Given this, it is plausible that the observed correlations in Sapap3-KOs between severity of these OCD-relevant phenotypes and reduced dopamine receptor density may reflect compensatory downregulation of receptors in response to increased dopamine tone. This has been proposed as a likely mechanisms for decreased striatal dopamine receptor binding observed using PET imaging in OCD patients^16^, and is supported by our observation of increased DAT binding density in Sapap3-KOs which may reflect increased dopaminergic innervation/release (Figure 4B). Interestingly, our prior study assessing the concentration of dopamine and its metabolites in *ex vivo* striatum using high performance liquid chromatography (HPLC) did not provide evidence of hyperdopaminergic tone in Sapap3-KO mice^46^. This suggests that measurement of behavior-associated dopamine release *in vivo* may be necessary to identify the changes in striatal dopamine suggested by our data. This possibility is in line with clinical studies demonstrating that cues associated with compulsive behavior are necessary to detect increased striatal dopamine in disorders associated with compulsive behavior^21-24^.

To our knowledge, these studies are the first to thoroughly investigate the longitudinal progressive development of the compulsive grooming phenotype in Sapap3-KOs, and raise the important question of whether there are distinct neural changes that lead to low vs high grooming KOs. Recently it has been reported that Sapap3-KO mice do not show age-dependent changes in grooming phenotype in a study using a cross-sectional design in a group of ∼2-5 month old Sapap3-KO mice^47^. In contrast, our within-subjects data tracking grooming from 2-8 months of age suggests that distinct trajectories for development of compulsive behavior exist among individual Sapap3-KOs. We show that, grooming progressively escalates to a point where it continues despite the presence of skin lesions in a subset of KOs, and that mice with lesions show significantly elevated grooming relative to non-lesioned KO mice (Figure 1D and S2F-H). Given the significant heterogeneity in compulsive grooming severity in Sapap3-KOs, observed both within animals (across time) and between animals at a given age, it will be important to report grooming severity in future research detailing the molecular, cellular, and circuit disturbances present in this model. Careful examination of these individual differences may help to understand neural mechanisms associated with resilience vs susceptibility to the development of compulsive behavior.

There are several limitations of these studies that will be important to address in future research. First, though there is some evidence that rare *Sapap3* variants play a role in OCD^48^, *Sapap3* is not a leading candidate for OCD genetic risk (though note that current genome wide association studies in OCD are underpowered^49^). However, our lab recently demonstrated decreased *Sapap3* gene expression in orbitofrontal cortex and striatum in *post mortem* tissue from human subjects with OCD, supporting a potential role for Sapap3 in OCD pathophysiology^50^. In further support of its use for OCD research, the Sapap3-KO mouse has abnormalities in OCD-relevant behavioral constructs (compulsive behavior, anxiety, executive dysfunction, and now disruption of sensorimotor gating), and exhibits therapeutic response to treatments that can be effective in OCD patients including repeated fluoxetine administration and DBS^32,51^. Nonetheless, further examination of the relationship between striatal dopamine, compulsive behavior, and sensorimotor gating is warranted in other OCD-relevant model systems and human subjects with OCD to better understand the relevance of our findings to OCD pathophysiology. Another limitation is that these studies rely on correlative evidence to link striatal dopamine receptor changes with severity of OCD-relevant behaviors. We propose that our observations provide indirect evidence that increased striatal dopamine is a common mechanism for generation of distinct OCD-relevant behaviors. We believe these findings warrant further causal mechanistic examination of dopaminergic regulation of behavioral disturbances relevant to OCD, which is beyond the scope of the current studies.

These studies are the first to demonstrate an association between the development of OCD-relevant compulsive grooming and disruption of PPI in a transgenic mouse model. Through *ex vivo* neurochemical analysis of brain tissue from mice that have undergone behavioral testing, we have identified a potential role of reduced NAcC dopamine receptor density in the manifestation of these maladaptive behaviors. By leveraging heterogeneity in the penetrance and progression of OCD-relevant behaviors in this preclinical model system, we have highlighted associations that may be relevant to neural changes occurring in distinct subsets of patients with OCD and related disorders.

## Methods

### Animals

Male and female Sapap3-KO and WT littermates were bred from heterozygous x heterozygous breeding pairs on a C57/BL6 background, and were derived from a colony initially established at MIT by Dr Guoping Feng^32^. Mice were housed in groups of 2-5 same-sex mice in individually ventilated cages, with *ad libitum* access to standard chow and drinking water, in 12:12 light/dark cycle. All behavioral testing was conducted in the light cycle. All experiments were approved by the Institutional Animal Care and Use Committee at the University of Pittsburgh in compliance with National Institutes of Health (NIH) guidelines for the care and use of laboratory animals.

### Prepulse Inhibition testing

PPI was tested using SR-LAB startle response system (San Diego Instruments, San Diego, CA) with methods similar to those previously described^9,27,52^, to examine the effects of three prepulse intensities (3, 6 and 12dB above background noise levels) on the magnitude of acoustic startle reflex elicited by 120dB pulse of noise. All mice were tested in “startle only” session (10 × 120dB pulse alone trials) prior to first PPI session for habituation to testing procedures. PPI is calculated as follows:

[(mean of 12 pulse alone trials [block2/3])-(mean of 10 prepulse + pulse trials)/ (mean of 12 pulse alone trials [block2/3])] x 100

Because these studies examined mice at an age where poor hearing could potentially impact PPI, mice were excluded from analysis if they exhibited very weak startle response to pulse alone trials relative to measurements made in the absence of a startling pulse (“no stimulus” trials, Figure S1D). The following criteria were used to exclude mice with weak startle reflex likely reflecting poor hearing: (average of “pulse alone” trial amplitudes in blocks 2&3) < (3 x average of “no stimulus” trial amplitudes).

### Grooming testing

Prior to first grooming test, mice were habituated to acrylic testing chambers (8 × 8 x 12 inches) for 2 consecutive days (20 mins/session). Mice were then videotaped for a 20 minute session, and grooming was manually scored offline by trained experimenters blind to genotype and PPI values.

### Lesion severity scoring

A trained experimenter blind to genotype and mouse behavioral scores rated each animal for evidence of skin lesions on the day of PPI testing. Mice that had regions on the face or body where skin was raw, inflamed, or bloody were classified in the “lesion” group. In line with institutional IACUC protocols, mice classified with lesions were treated using topical antibiotic as needed (Neomycin and Polymyxin B Sulfates, and Bacitracin Zinc Opthalmic Ointment; Akorn, Lake Forest, IL), with the exclusion of the day of and prior to behavioral testing.

### Autoradiography

Fresh frozen brains were obtained from mice assessed in PPI and grooming tests, and processed according to standard protocols to detect dopamine D1 and D2/3 receptor and DAT binding using specific radioligands^10,53,54^. Mice were sacrificed by cervical dislocation followed by rapid decapitation. Brains were then extracted, frozen immediately on dry ice, and stored at - 80°C. 20μm sections were cut on a cryostat (Lecia, Wetzlar, Germany) and stored on slides (Fisherbrand, Pittsburgh, PA) at −80°C until autoradiography experiments were performed.

Slides for total binding (TB) included tissue from 2 animals from different experimental groups, with 4 striatal sections collected every 200μm. Slides for non-specific binding (NSB) included tissue from 4 animals with 2 sections collected every 400μm. Position of subjects on the slides was counter-balanced across groups to account for fluid distribution during radioisotope incubation, and all slides for a given assay were incubated on the same day with the same binding solutions. Slides were thawed for 1 hour before preincubation in buffer (except for DAT, no preincubation). Following preincubation, slides were allowed to completely airdry for >1 hour before incubation in radioligand solutions. The following radioisotopes were used: [^3^H]-SCH-23390 for D1 binding (2nM), [^3^H]-Raclopride for D2/3 binding (5nM), [^3^H]-GBR-12935 for DAT binding (2nM) (Perkin Elmer, Boston, MA). To terminate radioligand incubation, slides were dipped and then washed in ice cold buffer (3 × 5 mins for D1 and D2/3, and 4 × 30 mins DAT). Following washes slides were dipped in ice cold dH_2_O and allowed to completely airdry before they were opposed to BioMax MR film (Kodak, Rochester, NY) with ^3^H standards (American Radiolabeled Chemicals, Inc., St Louis, MO). More details regarding reagents and conditions used for specific autoradiography experiments are included in Table S1; these were based on published protocols^53,54^. MR films were developed using MINI-MED 90 X ray film development system (AFP Manufacturing, Peachtree City, GA). Developed films were then scanned at 2400dpi (Perfection V550 Photo scanner, Epson, Suwa, Japan) and luminosity was measured using Photoshop (Adobe, San Jose, CA) bilaterally in each region of interest on TB and NSB slides. These values were converted to radioactivity (uCi/g) using a standard curve, and average NSB radioactivity was subtracted from average TB to give specific binding (SB) used for analysis. Figure S4 shows representative images for both TB and NSB for each assay.

### Cohort details

#### Experiment 1: Comparison of PPI and compulsive grooming severity

Male mice (n=14 WT, 18 KO) were tested at ∼8 months of age. Mice were assessed in the grooming test and PPI paradigm, and lesion severity was recorded. 2 WTs were excluded from PPI analysis due to poor hearing, leaving 12 WTs.

#### Experiment 2: Autoradiography

Brains from animals used in experiment 1 were used for autoradiography.

#### Experiment 3: Longitudinal characterization

From 2 months of age (before Sapap3-KOs show lesions) until 8 months, grooming was tested monthly, followed ∼3 days later by PPI testing [n=24 WT (12 male), 23 KO (10 males)]. Analysis of behavioral data separated by sex demonstrated no evidence for sex differences (Figure S5). Sexes were therefore combined for all analyses presented. Prior to the first test (but at no other time-points), mice were habituated for each paradigm as described above. Animals with %PPI >2 SD outside group average for that age were kept in the repeated measures longitudinal analysis (across 7 timepoints) but were excluded from analysis at 6-8 months of age as appropriate [n excluded: WT: 6 months=2, 7 months=3, 8 months=2; KO-NL: 6 months=1, 7 months=1, 8 months =1 (same mouse across all points); KO-L: none at any month].

#### Experiment 4: Early grooming phenotype

To examine grooming early in development, mice [n=17 WT (9 males), 17 KO (9 males)] were acclimatized twice to grooming chambers between postnatal day (PND) 18-20. Mice were weaned at PND21, tested the following day, and then tested weekly until week 8. Data for week 8 of this experiment has been published elsewhere^39^, and is reproduced with permission (Figure S3).

### Statistical analysis

Welch’s t-tests were used for analyses of data where the groups had unequal variance (e.g. PPI when KOs were analysed as one heterogeneous group, Figure 1A). Nonparametric t-tests were used for analyses of data that were not normally distributed (e.g. grooming when KOs were analysed as one heterogeneous population, Figure 1B). Repeated measures ANOVAs were used to compare the effects of group or genotype across time and different prepulse intensities; however, analysis of group differences examined average PPI across the three prepulse intensities. Pearson’s R was used to assess correlations between different measurements. T-tests, correlations, and 1 or 2-way ANOVAs were performed using GraphPad Prism (La Jolla, CA; version 8.2.1). Mice were excluded if PPI values were more than 2 standard deviations outside their group mean or PPI data indicated poor hearing (as defined above). Graphs show mean ± SEM for bar graphs and best-fit linear regression ± 95% confidence intervals.

## Acknowledgements

We would like to thank Ms Leela Ekambarapu and Mr Vinayak Banerjee for assistance with grooming assessments, Dr David Volk for use of his laboratory’s radioactive standards, and Dr Bianca Jupp for advice regarding autoradiography protocols. These studies were supported by BRAINS R01MH104255, McKnight Neuroscience Scholar Award, MQ Fellows Award, Burroughs Wellcome Fund CAMS, and Klingenstein Fellowship in the Neurosciences to S.E.A.

## Author Contributions

E.E.M. and S.E.A. conceptualized the studies and wrote the manuscript, E.E.M, A.Y.W. and A.S.W. performed behavioral experiments, E.E.M. and L.M.S. performed autoradiography experiments. All authors contributed to and approved the final manuscript.

## Additional Information

**Supplementary information** accompanies this paper

**Competing Interests**: The authors declare that they have no competing interests.

## Data Availability

The data are available from the corresponding author upon request.

## Supplementary materials

### Results

#### PPI and startle in Sapap3-KOs with and without lesions

%PPI increased with increasing prepulse intensity similarly in WT and KO mice (main effect PP, F_(2,56)_=87.4, p<0.0001; main effect genotype: p=0.14, PP x genotype interaction p=0.86; Figure S1A). Startle amplitude did not differ between the groups during the trial blocks used to calculate PPI (2 and 3) (main effect of group, p=0.43; Figure S1B). However, there was a trend for differences in startle habituation between the groups (Figure S1C, group x block interaction p=0.06). KO-L showed a trend for reduced startle in block 1 (vs KO-NL and WT, p<0.06, t_108_>2.3) and stable startle across the 4 blocks, whereas WT and KO-NL showed typical startle habituation. The amplitude of movement detected during trials when no stimulus was presented was reduced in both KO groups relative to WT (Figure S1D, F_(2,27)_=8.6, p=0.001). Bodyweight was also reduced in KOs relative to WT, although this effect was most pronounced in KO-L, which were significantly lighter than both WT and KO-NL (Figure S1E, F_(2,27)_=28.5, p<0.0001).

**Figure S1.**
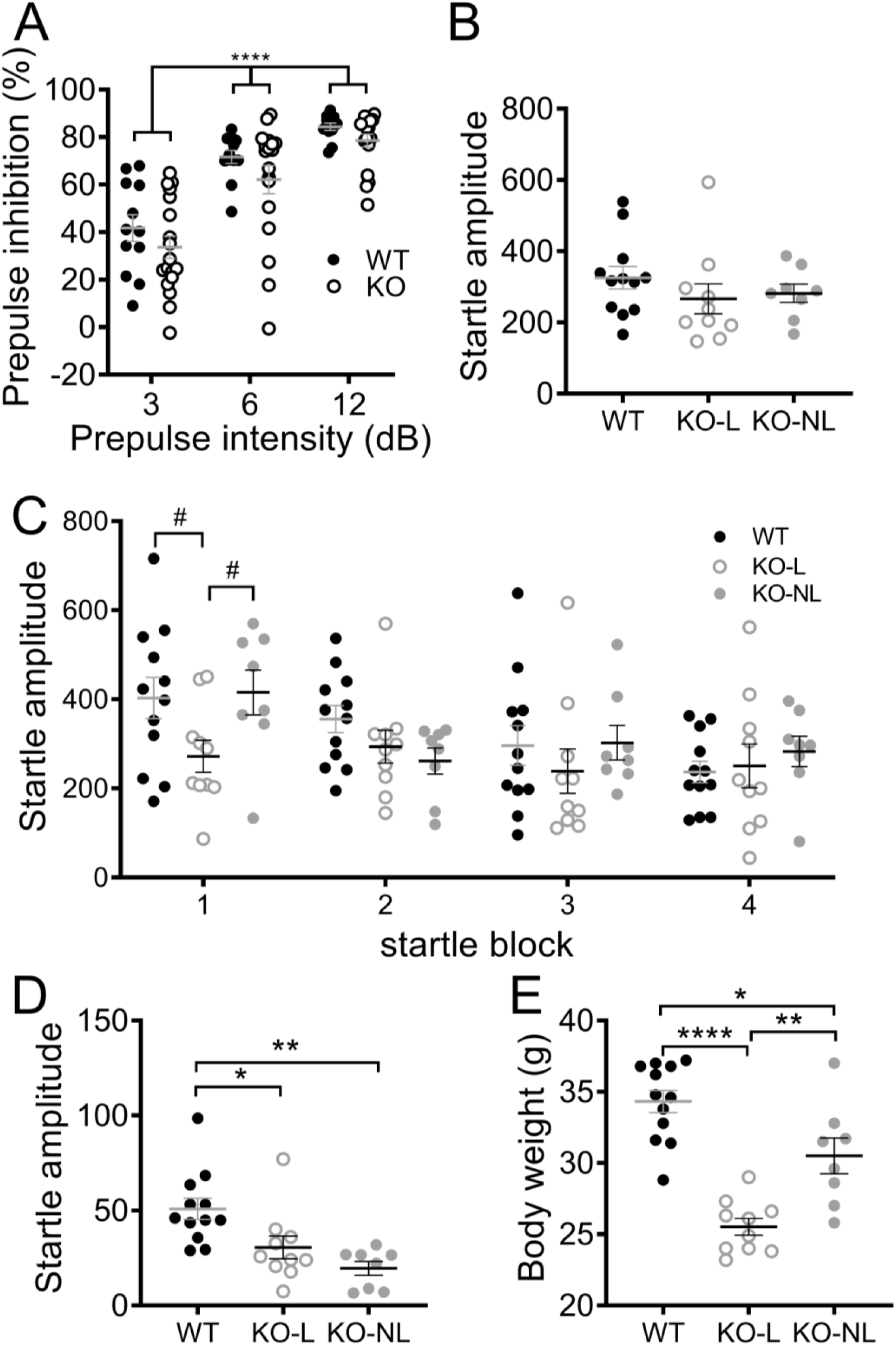
PPI and startle in Sapap3-KOs with and without lesions. A) PPI increases with increasing prepulse magnitude to a similar extent in WT and KO mice. B) Startle amplitude during 120dB pulse only trials in blocks 2/3 (used to calculate %PPI) does not differ between WT, KO-L and KO-NL. C) A trend was observed towards altered habituation to startling stimulus (120dB pulse only trials) between groups across 4 blocks of testing (p=0.058). Post-hoc tests demonstrated that this trend was driven by the KO-L group, which showed reduced startle during the first block relative to the other groups (t(108)>2.3, p<0.06), and an attenuation of habituation across the subsequent blocks relative to WT and KO-NL (block 1 vs block 4: KO-L, p>0.99; KO-NL t(81)=2.7, p=0.045; WT t(81)=4.2, p= 0004). D) Movement detected during “no stimulus” trials was reduced in KO-L and KO-NL relative to WT (KO-L vs WT t(27)=2.8, p=0.03, KO-NL vs WT t(27)=4.0, p=0.0014), but did not differ between KO-L and KO-NL (p=0.57). E) Body weight was reduced in KO-L and KO-NL relative to WT (KO-L vs WT t(27)=7.5, p<0.001, KO-NL vs WT t(27)=3.0, p=0.014). KO-L also showed lower body weight relative to KO-NL (t(27)=3.9, p=0.0019). In panel A, * indicates main effect of prepulse intensity, in panel C-E */# indicates results of Bonferroni post-hoc tests. **** p<0.0001; **p<0.01, *p<0.05, # 0.1>p>0.05. n=14 WT, 18 KO (10 KO-L, 8 KO-NL). KO= knockout, KO-L= KOs with lesions, KO-NL= KOs without lesions, WT= wild-type.

#### Longitudinal progression of changes in PPI, startle, and grooming in Sapap3-KO mice

PPI significantly increased across 2-8 months of age in both groups (Figure S2A; F_(6, 270)_=29.2, p<0.0001), and there was a trend for a reduction in PPI in Sapap3-KOs compared to WTs across 2-8 months of age (p=0.095). However, there were significant age-dependent differences in startle between the genotypes (Figure S2B; interaction p=0.001), with Sapap3-KOs showing reduced startle relative to WT between 2-5 months of age. Therefore, PPI could only be reliably compared between WT, KO-NL, and KO-L between 6-8 months of age when startle was comparable between the groups (Figure S2C-E; 1-way ANOVA comparing groups: 6 months p=0.13, 7 months p=0.65, 8 months p=0.18). Grooming was increased in KO-L relative to WT and KO-NL, and KO-NL relative to WT, from 6-8 months of age (Figure S2F-H).

**Figure S2.**
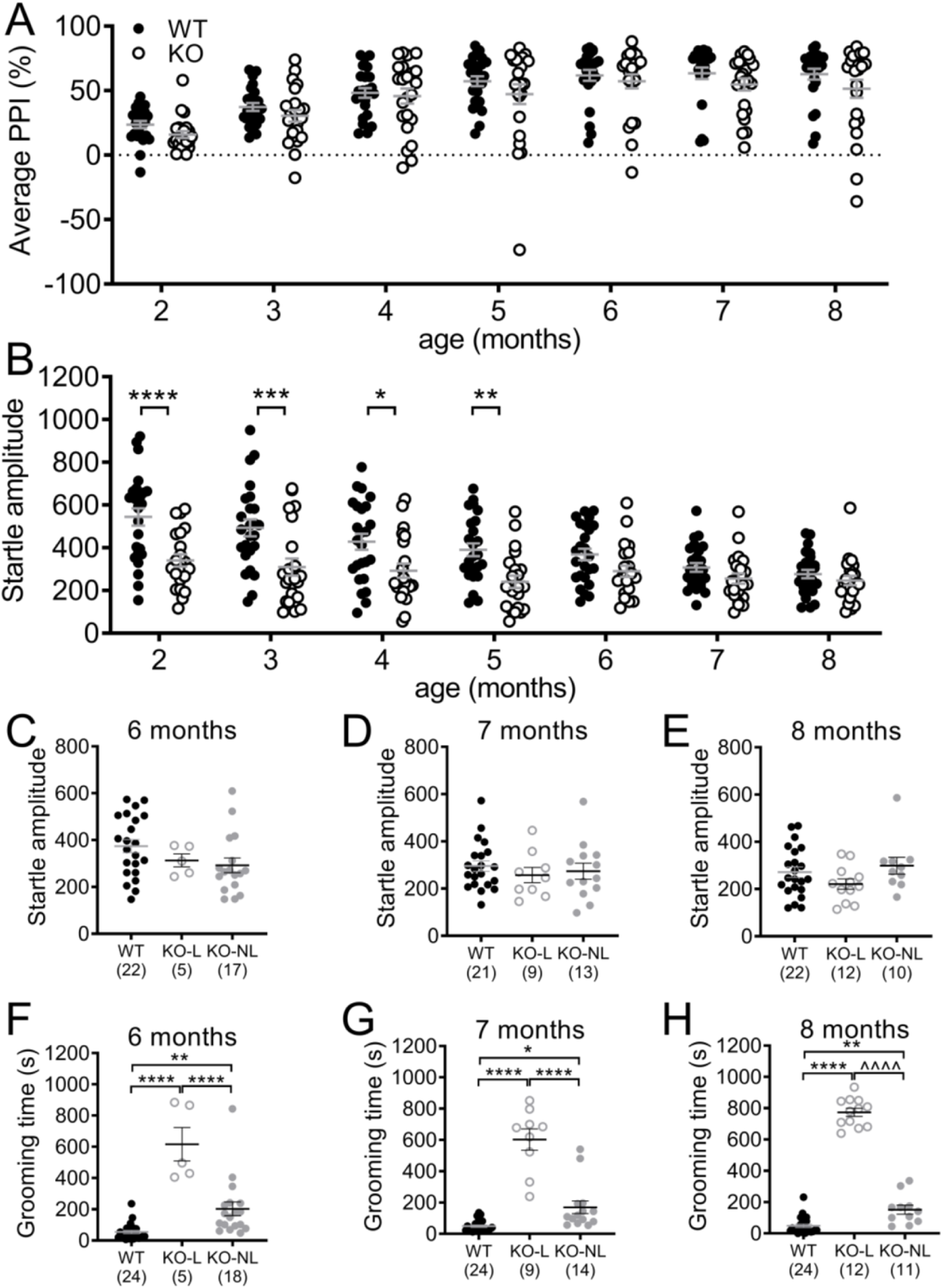
Longitudinal progression of changes in PPI, startle and grooming in Sapap3-KO mice. **A)** PPI significantly increased across 2-8 months of age in both groups. There was a trend for a reduction in PPI in Sapap3-KOs compared to WTs across 2-8 months of age (p=0.095). **B)** There were significant age-dependent differences in startle between the genotypes (interaction p=0.001), with Sapap3-KOs showing reduced startle relative to WT from 2-5 months of age. Therefore, PPI could only be reliably compared between WT, KO-NL and KO-L from 6-8 months of age when startle was comparable between the groups. Importantly, when KOs were separated into KO-L and KO-NL, startle was not significantly different from WT between 6-8 months of age **(C-E)**. Grooming was increased in KO-L relative to WT and KO-NL, and KO-NL relative to WT, from 6-8 months of age **(F-H)**. Note, for panels A,B, F-H all mice are included (n=24 WT, 23 KO), whereas for panels C-E mice excluded from PPI analysis have been removed, resulting in different group sizes. KO= knockout, KO-L= KOs with lesions, KO-NL= KOs without lesions, WT= wild-type, PPI= prepulse inhibition.

#### Early grooming phenotype

Assessment of post-weaning grooming phenotype from 3-8 weeks of age revealed a significant age dependent difference in grooming in Sapap3-KOs (Figure S3; age x genotype interaction, F_(5,160)_=8.5, p<0.0001). Post-hoc tests revealed that grooming was increased in WT relative to KO at 3 weeks of age, whereas grooming was increased in KO relative to WT at 7-8 weeks of age.

**Figure S3.**
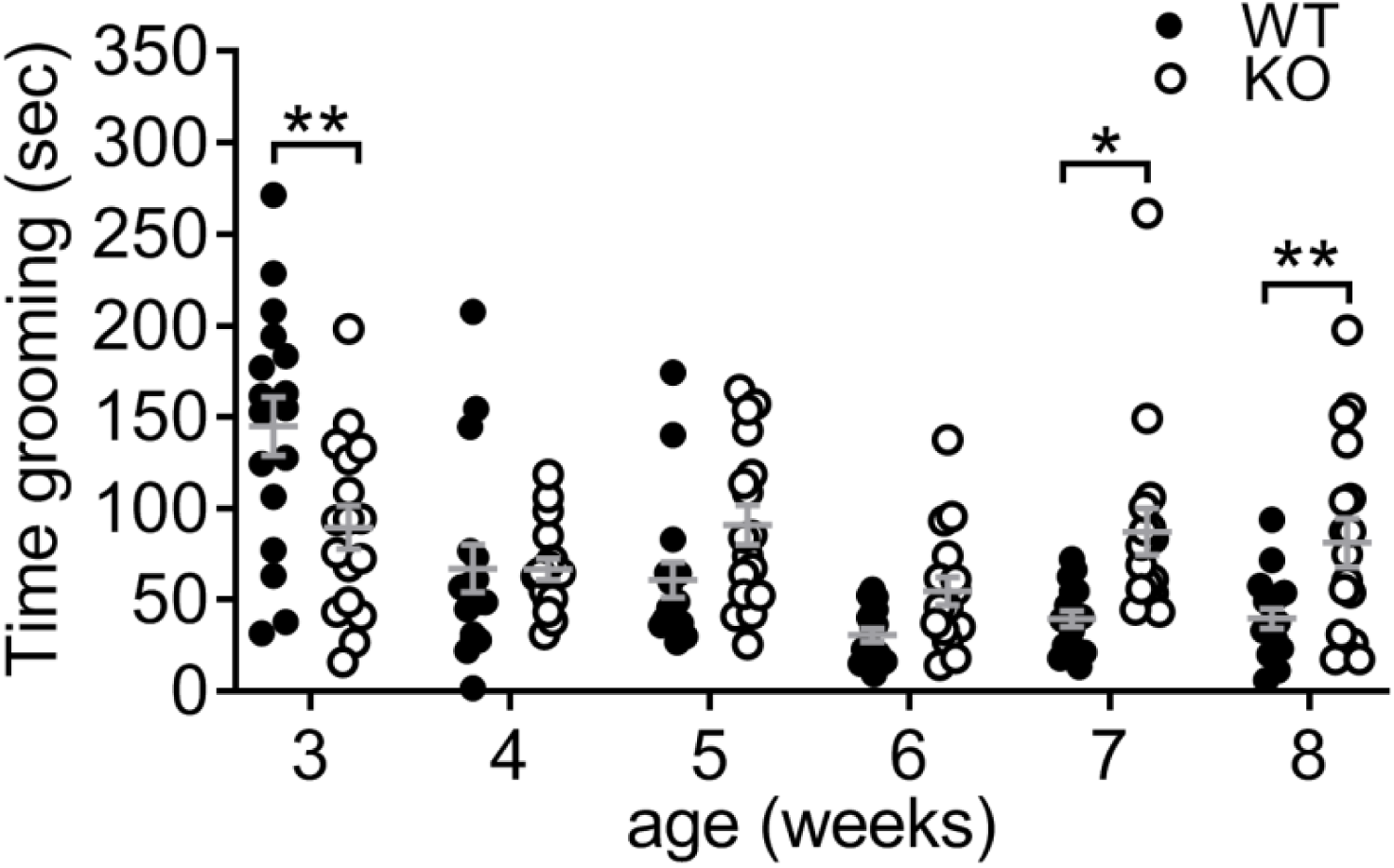
Early progression of grooming phenotype in Sapap3-KO mice. The early progression of the grooming phenotype was examined during the first month of life after weaning (3-8 weeks of age). Data from 8 weeks has been published elsewhere^1^. There were time-dependent differences in grooming between the genotypes. Post-hoc tests demonstrated that WTs showed elevated grooming at 3 weeks of age relative to KOs (t_(192)_=3.8, p=0.0011), whereas at 7 and 8 weeks of age grooming was elevated in Sapap3-KOs (7 weeks, t_(192)_=3.3, p=0.007; 8 weeks, t_(192)_=2.9, p=0.029). * indicates results of Bonferroni post-hoc test, * p<0.05, **p<0.01. n=17 WT (9 males), 15 KO (7 males). KO = knockout, WT= wild-type mice.

#### Representative images of total binding and non-specific binding sections for autoradiography assays

Receptor binding was calculated by measuring luminosity in striatal regions of interest (Figure 4SB) on total binding (TB) and nonspecific binding (NSB) slides (Figure S4A for representative images). These values were then converted to radioactivity (uCi/g) using a tritium standard curve, and NSB radioactivity was subtracted from TB radioactivity to give specific receptor/transporter binding value.

**Figure S4.**
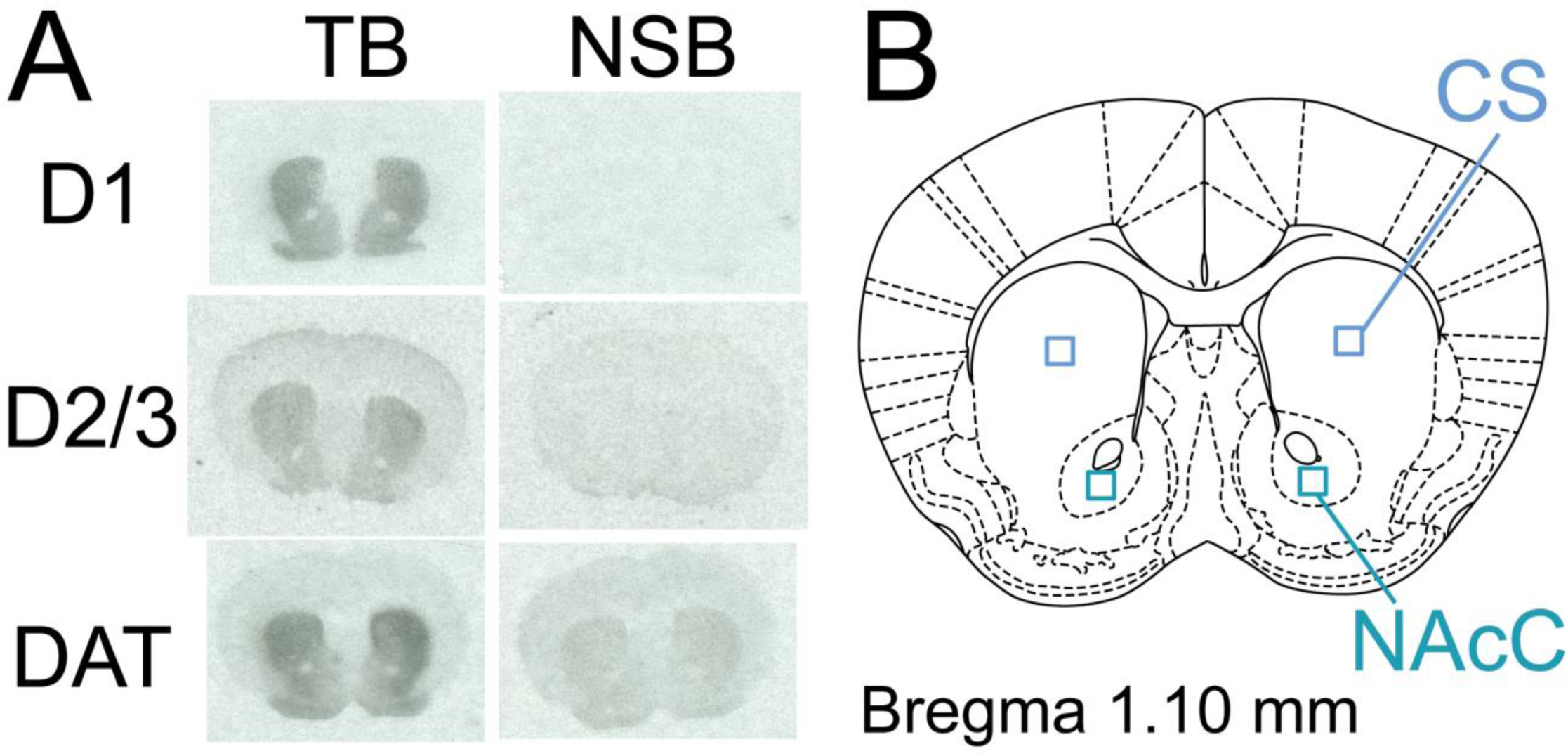
Representative autoradiography images and regions of interest. **A)** Representative TB and adjacent NSB autoradiographs from striatum for D1 receptor, D2/3 receptor, and DAT binding. **B)** Schematic diagram of coronal section with positioning of regions of interest used for quantification. CS= central striatum, NAcC= Nucleus accumbens core, DAT= dopamine transporter, TB= total binding, NSB= non-specific binding.

#### No sex differences in disruption of PPI or compulsive grooming in Sapap3-KO mice

Male and female mice were used in experiment 3. The data from these studies were used to investigate sex differences, given recent observations that PPI was disrupted in female OCD patients relative to healthy controls, but not in males^2^. In experiment 3, at 8 months of age there were no effects of sex (Figure S2C, main effect of group F_(2,38)_=6.6, p=0.003, main effect of sex, p=0.71, sex x group interaction p=0.82). Planned Bonferroni post-hoc tests were performed to compare the groups in each sex separately, and although this analysis was underpowered it supported disruption of PPI in KO-L of both sexes (Figure S5A: male KO-L vs WT, t_(38)_=2.2, p=0.0921, KO-L vs KO-NL t_(38)_=2.5, p=0.045; female KO-L vs WT, t_(38)_=2.5, p=0.05; for all other comparisons p>0.20). Severity of compulsive grooming phenotype also did not differ between the sexes (Figure S5B; main effect of group, p<0.0001; main effect of sex, p=0.948, interaction p=0.509).

**Figure S5.**
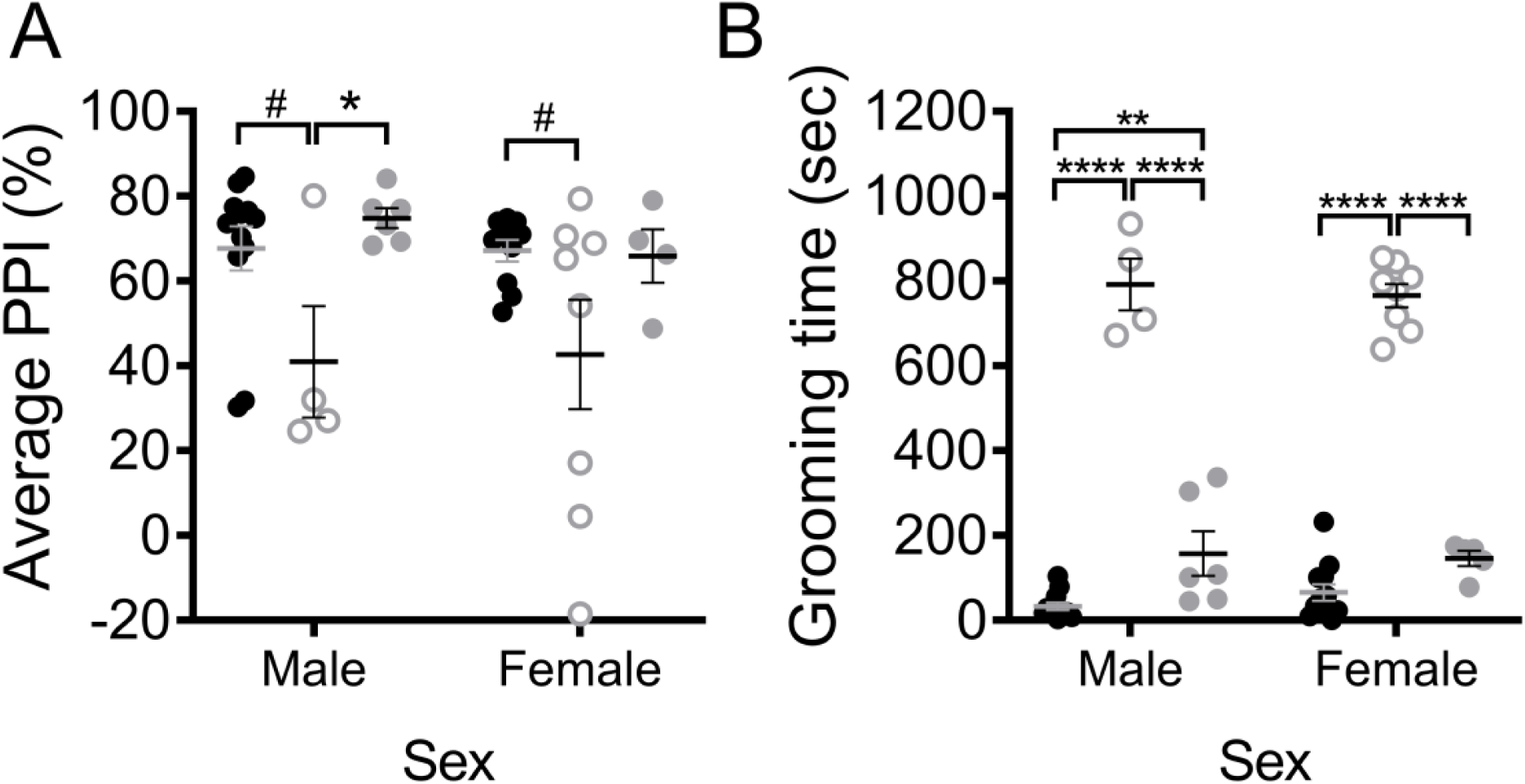
No sex differences in disruption of PPI or compulsive grooming in Sapap3-KO mice. **A)** Data from 8 month old mice from the longitudinal testing cohort (experiment 3) showed no evidence for sex differences in disruption of PPI. **B)** Compulsive grooming severity from experiment 3 also showed no evidence for sex differences. * indicates significant Bonferroni post-hoc test. # 0.1>p>0.05; * p<0.05,** p<0.01, ****p<0.0001. male: n=4 KO-L, 6KO-NL, 12 WT; female: n= 8 KO-L, 4 KO-NL, 10 WT. KO-L= KO mice with lesions, KO-NL= knockout mice without lesions, WT= wild-type, PPI= prepulse inhibition.

### Discussion

#### Detection of early changes in PPI and grooming in KOs

PPI could not be reliably compared between Sapap3-KOs and WT at early ages due to differences in startle amplitude. Therefore future studies may seek to “titrate” startle intensity between the groups (i.e. use lower intensity acoustic stimulus to induce startle in WT mice) so that startle is similar between the groups, allowing for more reliable comparison of PPI at earlier ages during the emergence of compulsive grooming in Sapap3-KOs^3^.

In the longitudinal study (experiment 3), we were unable to replicate the findings of experiment 4 demonstrating an increase in grooming in Sapap3-KO mice at 8 weeks/2 months of age. Instead, we first saw grooming-related genotype differences emerge at 5 months of age. Detection of the earliest, mildest grooming phenotype may therefore be dependent on regular testing and handling, which may be the cause of lower variance and lower mean levels of grooming in WT control mice in experiment 4 (weekly testing during experiment 4: 8 weeks of age WT mean = 39.7, SD= 22.5; monthly testing during experiment 3: 2 months of age WT mean= 67.0, SD= 47.4).

**Table S1:** reagents and conditions used for autoradiography.

| Receptor | Ligand | Ligand concentrations |  | Binding conditions |  |  |  | NSB | Exposure time |
| --- | --- | --- | --- | --- | --- | --- | --- | --- | --- |
|  |  | M | Ci/mmol | Temp | Time | Inhibitors | Buffer |  |  |
| D1 | [<sup>3</sup>H]-SCH 23390<sup>1</sup> | 2nM | 81.9 | RT | 1 hr | 1μM Ketanserin<sup>2</sup> | 50mM Tris-HCl<sup>2</sup>, 1mM MgCl<sub>2</sub><sup>2</sup>, 120mM NaCl<sup>2</sup>, 2mM CaCl<sub>2</sub><sup>2</sup>, 5mM KCl<sup>2</sup> | 10μM cis-flupenthixol<sup>2</sup> | 3.5 weeks |
| D2/3 | [<sup>3</sup>H]-Raclopride<sup>1</sup> | 5nM | 73.8 | RT | 1.5 hr | None | Same as D1 | 10μM Haloperidol<sup>2</sup> | 7.5 weeks |
| DAT | [<sup>3</sup>H]-GBR 12935<sup>1</sup> | 2nM | 41.3 | 4°C | 20 hrs | 1μM cis-Flupenthixol<sup>2</sup> | 50mM HNa<sub>2</sub>PO<sub>4</sub><sup>2</sup>, 70mM NaCl<sup>2</sup>, 0.025% BSA<sup>2</sup> | 10μM Mazindol<sup>2</sup> | 3 weeks |

Superscript indicates chemical manufacturer, with <sup>1</sup>= Perkin Elmer (Boston, MA), <sup>2</sup>= Sigma Aldrich (St Louis, MO), BSA= Bovine Serum Albumin, RT = Room temperature (~25°C), DAT= dopamine transporter.

