## Supplementary materials for "Disruption of prepulse inhibition is associated with severity of compulsive behavior and nucleus accumbens dopamine receptor changes in Sapap3 knockout mice"

#### Results

##### PPI and startle in Sapap3-KOs with and without lesions

%PPI increased with increasing prepulse intensity similarly in WT and KO mice (main effect PP, F_(2,56)_=87.4, p<0.0001; main effect genotype: p=0.14, PP x genotype interaction p=0.86; Figure S1A). Startle amplitude did not differ between the groups during the trial blocks used to calculate PPI (2 and 3) (main effect of group, p=0.43; Figure S1B). However, there was a trend for differences in startle habituation between the groups (Figure S1C, group x block interaction p=0.06). KO-L showed a trend for reduced startle in block 1 (vs KO-NL and WT, p<0.06, t_108_>2.3) and stable startle across the 4 blocks, whereas WT and KO-NL showed typical startle habituation. The amplitude of movement detected during trials when no stimulus was presented was reduced in both KO groups relative to WT (Figure S1D, F_(2,27)_=8.6, p=0.001). Bodyweight was also reduced in KOs relative to WT, although this effect was most pronounced in KO-L,
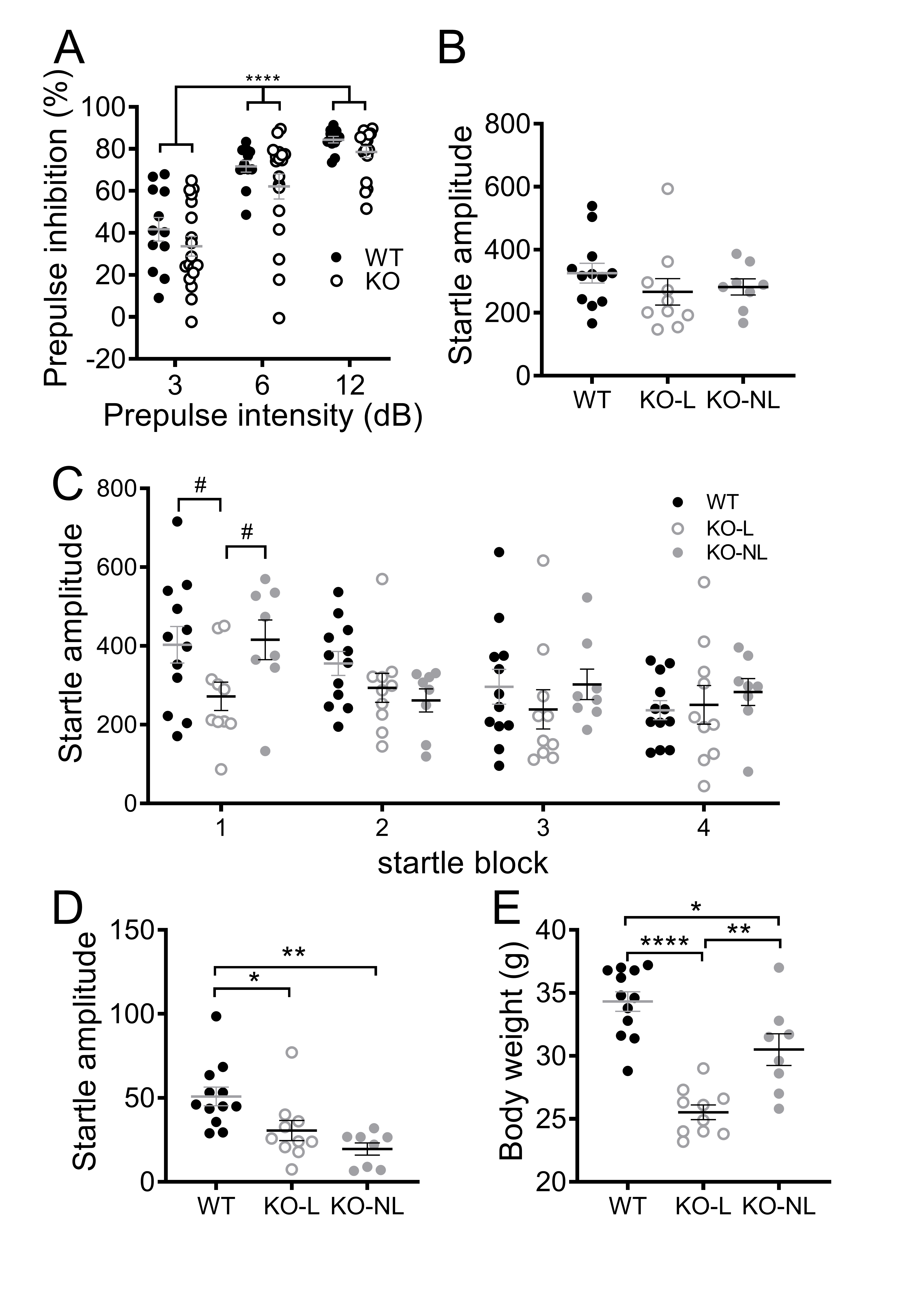
which were significantly lighter than both WT and KO-NL (Figure S1E, F_(2,27)_=28.5, p<0.0001).

###### **Figure S1.** PPI and startle in Sapap3-KOs with and without lesions. **A)** PPI increases with increasing prepulse magnitude to a similar extent in WT and KO mice. **B)** Startle amplitude during 120dB pulse only trials in blocks 2/3 (used to calculate %PPI) does not differ between WT, KO-L and KO-NL. **C)** A trend was observed towards altered habituation to startling stimulus (120dB pulse only trials) between groups across 4 blocks of testing (p=0.058). Post-hoc tests demonstrated that this trend was driven by the KO-L group, which showed reduced startle during the first block relative to the other groups (t_(108)_>2.3, p<0.06), and an attenuation of habituation across the subsequent blocks relative to WT and KO-NL (block 1 vs block 4: KO-L, p>0.99; KO-NL t_(81)_=2.7, p=0.045; WT t_(81)_=4.2, p= 0004). **D)** Movement detected during “no stimulus” trials was reduced in KO-L and KO-NL relative to WT (KO-L vs WT t_(27)_=2.8, p=0.03, KO-NL vs WT t_(27)_=4.0, p=0.0014), but did not differ between KO-L and KO-NL (p=0.57). **E)** Body weight was reduced in KO-L and KO-NL relative to WT (KO-L vs WT t_(27)_=7.5, p<0.001, KO-NL vs WT t_(27)_=3.0, p=0.014). KO-L also showed lower body weight relative to KO-NL (t_(27)_=3.9, p=0.0019). In panel A, * indicates main effect of prepulse intensity, in panel C-E */# indicates results of Bonferroni post-hoc tests. **** p<0.0001; **p<0.01, *p<0.05, # 0.1>p>0.05. n=14 WT, 18 KO (10 KO-L, 8 KO-NL). KO= knockout, KO-L= KOs with lesions, KO-NL= KOs without lesions, WT= wild-type.

##### Longitudinal progression of changes in PPI, startle, and grooming in Sapap3-KO mice

PPI significantly increased across 2-8 months of age in both groups (Figure S2A; F_(6, 270)_=29.2, p<0.0001), and there was a trend for a reduction in PPI in Sapap3-KOs compared to WTs across 2-8 months of age (p=0.095). However, there were significant age-dependent differences in startle between the genotypes (Figure S2B; interaction p=0.001), with Sapap3-KOs showing reduced startle relative to WT between 2-5 months of age. Therefore, PPI could only be reliably compared between WT, KO-NL, and KO-L between 6-8 months of age when startle was comparable between the groups (Figure S2C-E; 1-way ANOVA comparing groups: 6 months p=0.13, 7 months p=0.65, 8 months p=0.18). Grooming was increased in KO-L relative
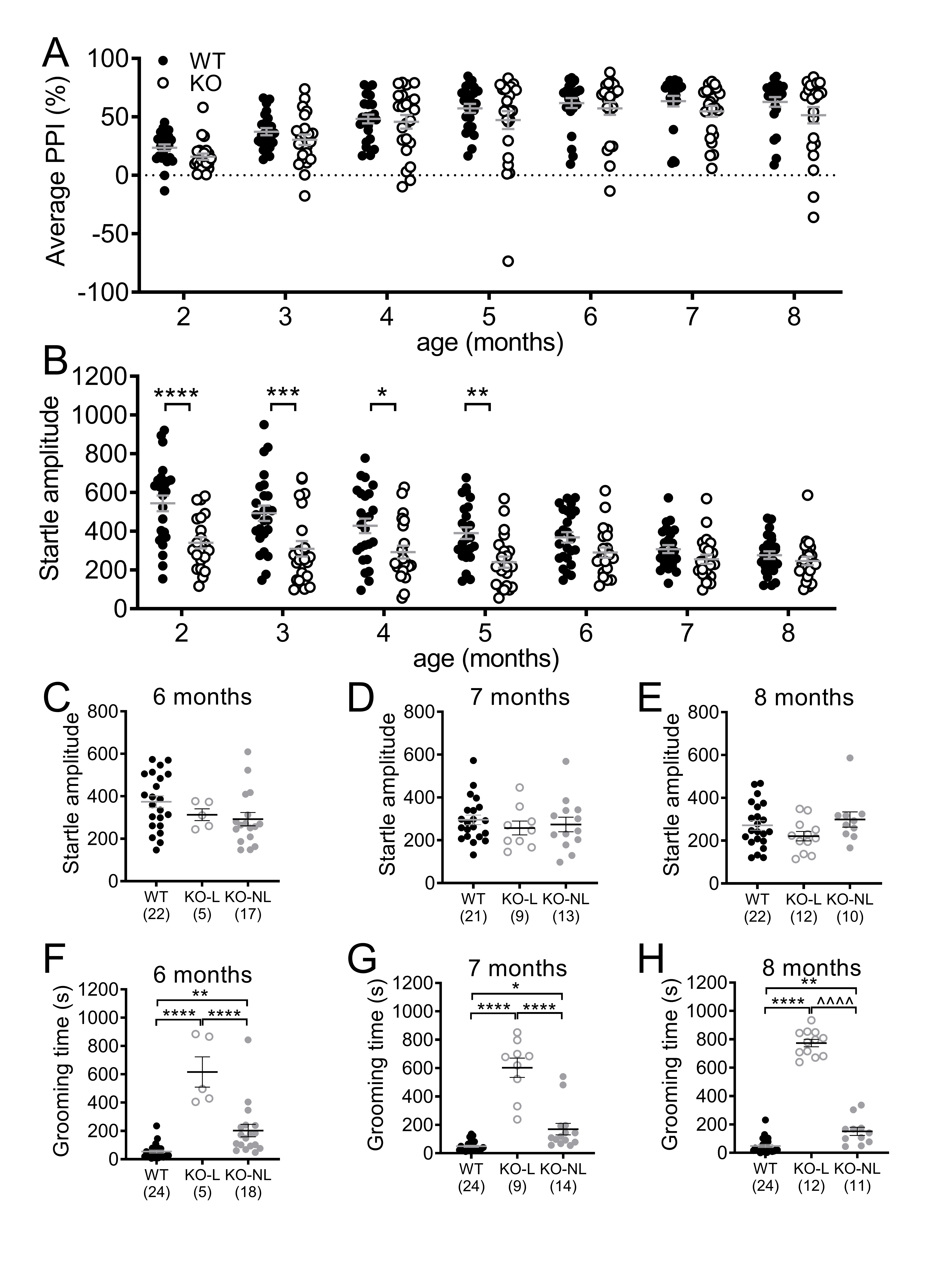
to WT and KO-NL, and KO-NL relative to WT, from 6-8 months of age (Figure S2F-H).

###### **Figure S2.** Longitudinal progression of changes in PPI, startle and grooming in Sapap3-KO mice. **A)** PPI significantly increased across 2-8 months of age in both groups. There was a trend for a reduction in PPI in Sapap3-KOs compared to WTs across 2-8 months of age (p=0.095). **B)** There were significant age-dependent differences in startle between the genotypes (interaction p=0.001), with Sapap3-KOs showing reduced startle relative to WT from 2-5 months of age. Therefore, PPI could only be reliably compared between WT, KO-NL and KO-L from 6-8 months of age when startle was comparable between the groups. Importantly, when KOs were separated into KO-L and KO-NL, startle was not significantly different from WT between 6-8 months of age **(C-E)**. Grooming was increased in KO-L relative to WT and KO-NL, and KO-NL relative to WT, from 6-8 months of age **(F-H)**. Note, for panels A,B, F-H all mice are included (n=24 WT, 23 KO), whereas for panels C-E mice excluded from PPI analysis have been removed, resulting in different group sizes. KO= knockout, KO-L= KOs with lesions, KO-NL= KOs without lesions, WT= wild-type, PPI= prepulse inhibition.

##### Early grooming phenotype

Assessment of post-weaning grooming phenotype from 3-8 weeks of age revealed a significant age dependent difference in grooming in Sapap3-KOs (Figure S3; age x genotype interaction, F_(5,160)_=8.5, p<0.0001). Post-hoc tests revealed that grooming was increased in WT relative to KO at 3 weeks of age, whereas grooming was increased in KO relative to WT at 7-8 weeks of age.

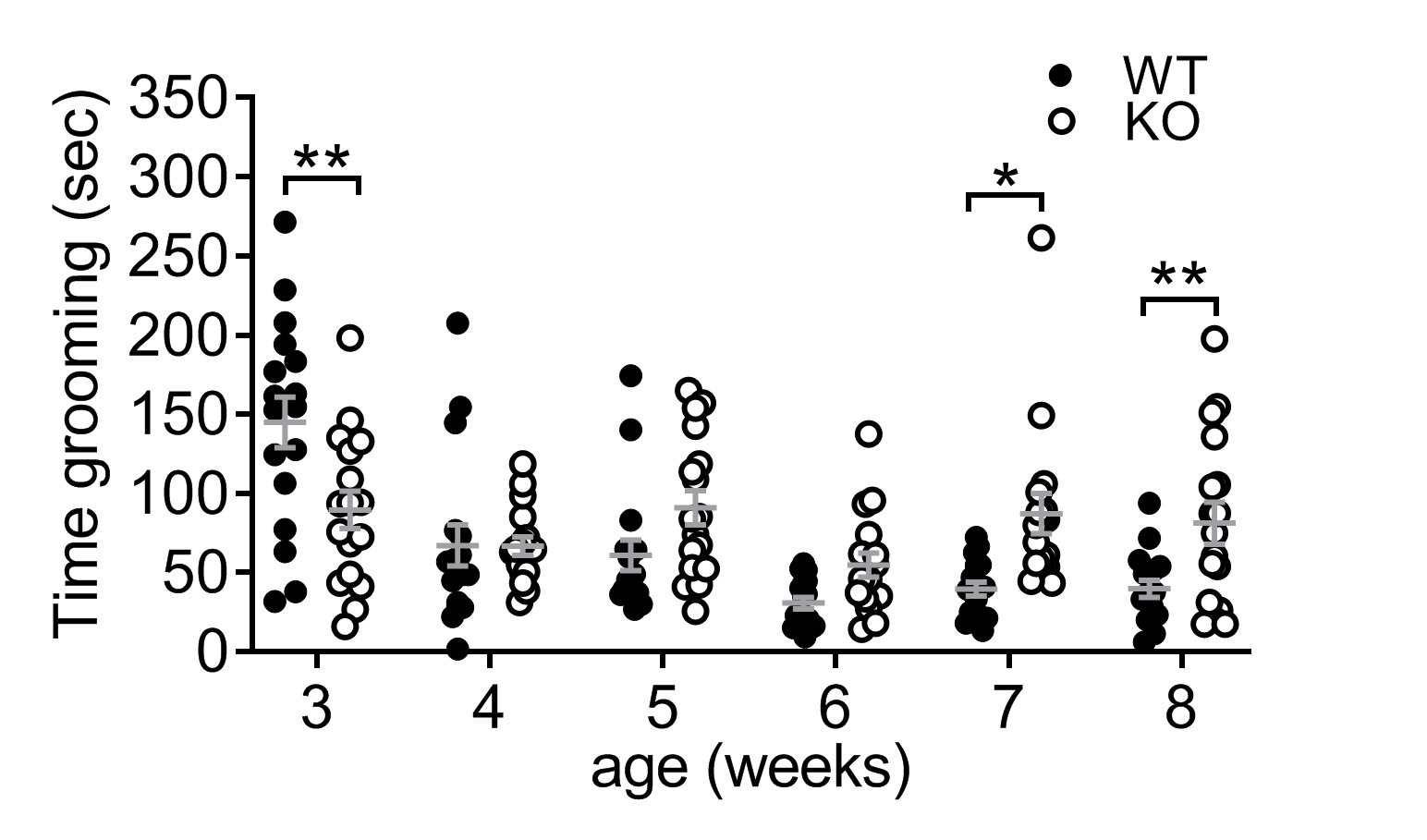

###### **Figure S3.** Early progression of grooming phenotype in Sapap3-KO mice. The early progression of the grooming phenotype was examined during the first month of life after weaning (3-8 weeks of age). Data from 8 weeks has been published elsewhere^1^. There were time-dependent differences in grooming between the genotypes. Post-hoc tests demonstrated that WTs showed elevated grooming at 3 weeks of age relative to KOs (t_(192)_=3.8, p=0.0011), whereas at 7 and 8 weeks of age grooming was elevated in Sapap3-KOs (7 weeks, t_(192)_=3.3, p=0.007; 8 weeks, t_(192)_=2.9, p=0.029). * indicates results of Bonferroni post-hoc test, * p<0.05, **p<0.01. n=17 WT (9 males), 15 KO (7 males). KO = knockout, WT= wild-type mice.

##### Representative images of total binding and non-specific binding sections for autoradiography assays

Receptor binding was calculated by measuring luminosity in striatal regions of interest (Figure 4SB) on total binding (TB) and nonspecific binding (NSB) slides (Figure S4A for representative images). These values were then converted to radioactivity (uCi/g) using a tritium standard curve, and NSB radioactivity was subtracted from TB radioactivity to give specific receptor/transporter binding value.

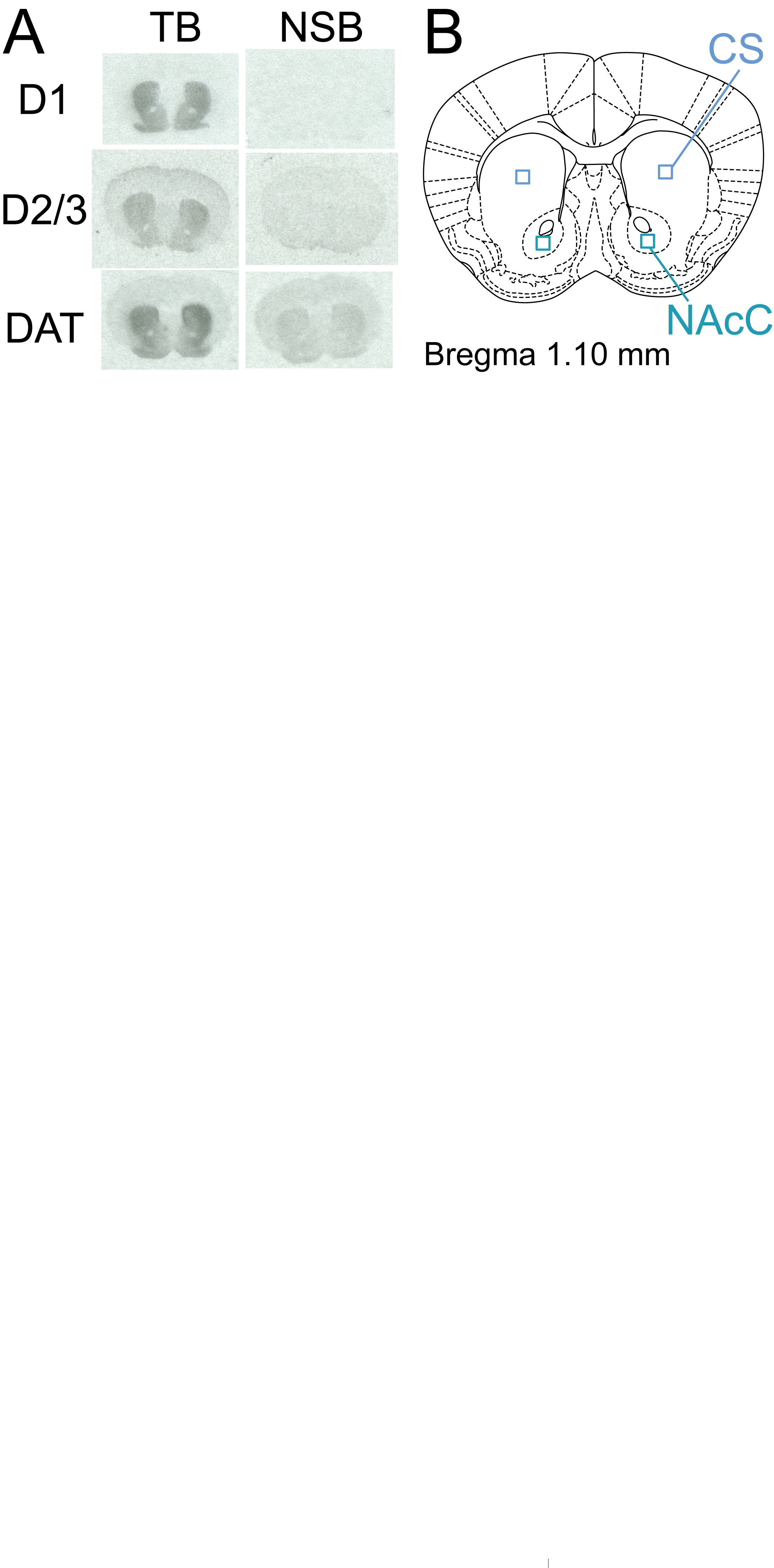

###### **Figure S4.** Representative autoradiography images and regions of interest. **A)** Representative TB and adjacent NSB autoradiographs from striatum for D1 receptor, D2/3 receptor, and DAT binding. **B)** Schematic diagram of coronal section with positioning of regions of interest used for quantification. CS= central striatum, NAcC= Nucleus accumbens core, DAT= dopamine transporter, TB= total binding, NSB= non-specific binding.

##### No sex differences in disruption of PPI or compulsive grooming in Sapap3-KO mice

Male and female mice were used in experiment 3. The data from these studies were used to investigate sex differences, given recent observations that PPI was disrupted in female OCD patients relative to healthy controls, but not in males^2^. In experiment 3, at 8 months of age there were no effects of sex (Figure S2C, main effect of group F_(2,38)_=6.6, p=0.003, main effect of sex, p=0.71, sex x group interaction p=0.82). Planned Bonferroni post-hoc tests were performed to compare the groups in each sex separately, and although this analysis was underpowered it supported disruption of PPI in KO-L of both sexes (Figure S5A: male KO-L vs WT, t_(38)_=2.2, p=0.0921, KO-L vs KO-NL t_(38)_=2.5, p=0.045; female KO-L vs WT, t_(38)_=2.5, p=0.05; for all other comparisons p>0.20). Severity of compulsive grooming phenotype also did not differ between the sexes (Figure S5B; main effect of group, p<0.0001; main effect of sex, p=0.948, interaction p=0.509).

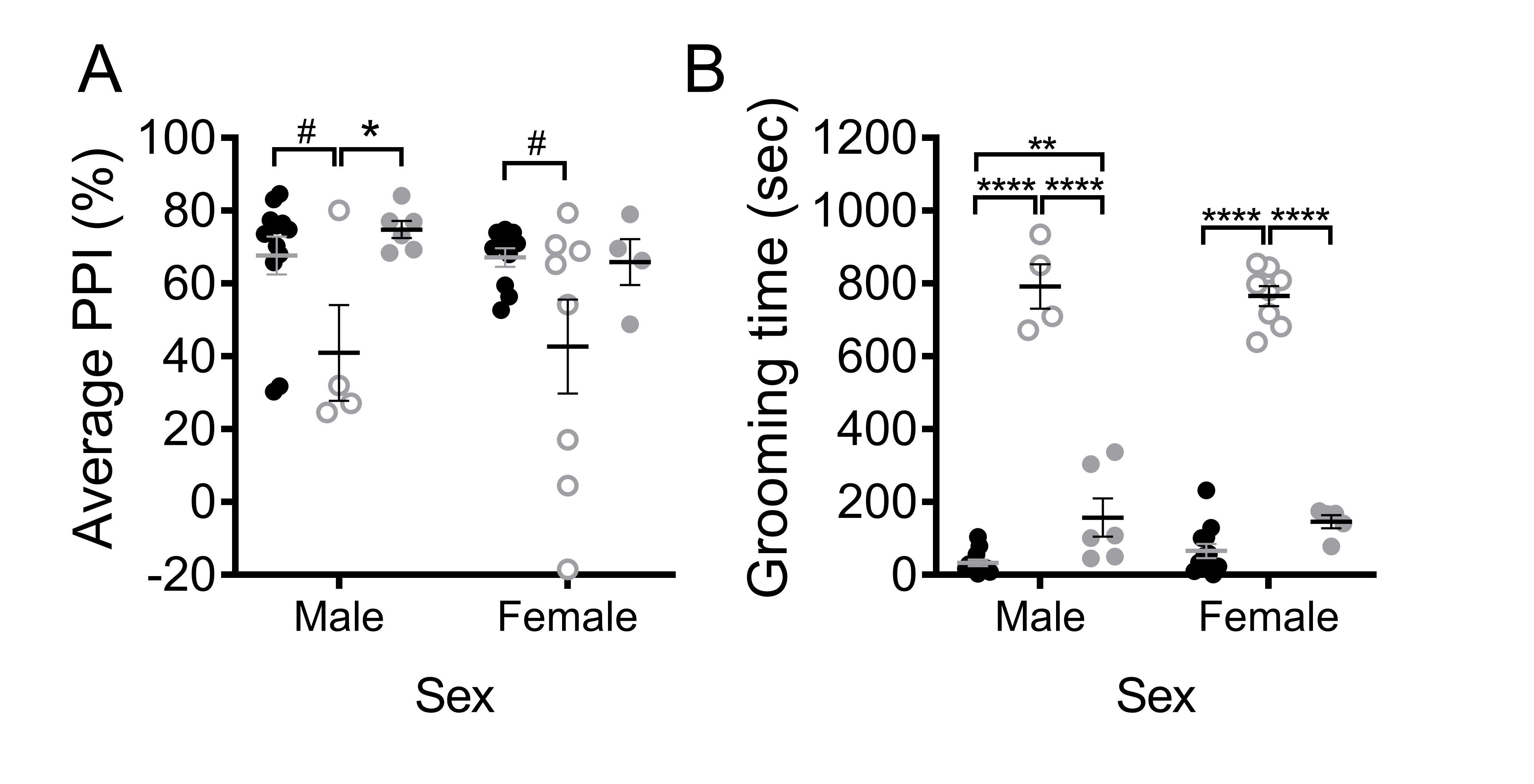

###### Figure S5. No sex differences in disruption of PPI or compulsive grooming in Sapap3-KO mice. **A)** Data from 8 month old mice from the longitudinal testing cohort (experiment 3) showed no evidence for sex differences in disruption of PPI. **B)** Compulsive grooming severity from experiment 3 also showed no evidence for sex differences. * indicates significant Bonferroni post-hoc test. # 0.1>p>0.05; * p<0.05,** p<0.01, ****p<0.0001. male: n=4 KO-L, 6KO-NL, 12 WT; female: n= 8 KO-L, 4 KO-NL, 10 WT. KO-L= KO mice with lesions, KO-NL= knockout mice without lesions, WT= wild-type, PPI= prepulse inhibition.

### Discussion

#### Detection of early changes in PPI and grooming in KOs

PPI could not be reliably compared between Sapap3-KOs and WT at early ages due to differences in startle amplitude. Therefore future studies may seek to “titrate” startle intensity between the groups (i.e. use lower intensity acoustic stimulus to induce startle in WT mice) so that startle is similar between the groups, allowing for more reliable comparison of PPI at earlier ages during the emergence of compulsive grooming in Sapap3-KOs^3^.

In the longitudinal study (experiment 3), we were unable to replicate the findings of experiment 4 demonstrating an increase in grooming in Sapap3-KO mice at 8 weeks/2 months of age. Instead, we first saw grooming-related genotype differences emerge at 5 months of age. Detection of the earliest, mildest grooming phenotype may therefore be dependent on regular testing and handling, which may be the cause of lower variance and lower mean levels of grooming in WT control mice in experiment 4 (weekly testing during experiment 4: 8 weeks of age WT mean = 39.7, SD= 22.5; monthly testing during experiment 3: 2 months of age WT mean= 67.0, SD= 47.4).

##### Table S1: reagents and conditions used for autoradiography

|  |  | Ligand concentrations | | Binding conditions | | | |  |  |
| --- | --- | --- | --- | --- | --- | --- | --- | --- | --- |
| Receptor | Ligand | M | Ci/mmol | Temp | Time | Inhibitors | Buffer | NSB | Exposure time |
| D1 | [^3^H]-SCH 23390**^1^** | 2nM | 81.9 | RT | 1 hr | 1µM Ketanserin**^2^** | 50mM Tris-HCl**^2^**, 1mM MgCl_2_**^2^**_,_ 120mM NaCl**^2^**, 2mM CaCl_2_**^2^**, 5mM KCl**^2^** | 10µM cis-flupenthixol**^2^** | 3.5 weeks |
| D2/3 | [^3^H]-Raclopride**^1^** | 5nM | 73.8 | RT | 1.5 hr | None | Same as D1 | 10µM Haloperidol**^2^** | 7.5 weeks |
| DAT | [^3^H]-GBR 12935**^1^** | 2nM | 41.3 | 4ºC | 20 hrs | 1µM cis-Flupenthixol**^2^** | 50mM HNa_2_PO_4_**^2^**, 70mM NaCl**^2^**, 0.025% BSA**^2^** | 10µM Mazindol**^2^** | 3 weeks |

###### Superscript indicates chemical manufacturer, with **^1^**= Perkin Elmer (Boston, MA), **^2^**= Sigma Aldrich (St Louis, MO), BSA= Bovine Serum Albumin, RT = Room temperature (~25ºC), DAT= dopamine transporter.
